# Emerging cooperativity between Oct4 and Sox2 governs the pluripotency network in early mouse embryos

**DOI:** 10.1101/2023.10.18.562912

**Authors:** Yanlin Hou, Zhengwen Nie, Qi Jiang, Sergiy Velychko, Sandra Heising, Ivan Bedzhov, Guangming Wu, Kenjiro Adachi, Hans R. Schöler

## Abstract

During the first lineage segregation, mammalian embryos generate the inner cell mass (ICM) and trophectoderm (TE). ICM gives rise to the epiblast (EPI) that forms all cell types of the body, an ability referred to as pluripotency. The molecular mechanisms that induce pluripotency in embryos remain incompletely elucidated. Using knockout (KO) mouse models in conjunction with low-input ATAC-seq and RNA-seq, we found that Oct4 and Sox2 gradually come into play in the early ICM, coinciding with the initiation of Sox2 expression. Oct4 and Sox2 activate the pluripotency-related genes through the putative OCT-SOX enhancers in the early ICM. Furthermore, we observed a substantial reorganization of chromatin landscape and transcriptome from the morula to the early ICM stages, which was partially driven by Oct4 and Sox2, highlighting their pivotal role in promoting the developmental trajectory towards the ICM. Our study provides new insights into the establishment of the pluripotency network in mouse preimplantation embryos.

## Introduction

Following fertilization, mouse embryos undergo a series of cell divisions and differentiation in the first few days. Zygotes and 2-cell blastomeres are totipotent and can form both embryonic and extra-embryonic supporting tissues. After several rounds of divisions, embryos undergo compaction at the morula stage and specifies the trophectoderm (TE) and inner cell mass (ICM). The ICM further segregates into the primitive endoderm (PE) and the epiblast (EPI) in the subsequent blastocyst stage. All cell types of the fetus can develop from the EPI, a feature known as pluripotency. During the establishment of pluripotency, dramatic molecular changes occur such as shifts in epigenetic modifications, chromatin accessibility and transcriptome (Deng *et al*, 2014; Guo *et al*, 2010; Wang *et al*, 2018; Wu *et al*, 2016; Zhang *et al*, 2016; Zhang *et al*, 2018). The precise mechanisms involved in establishing the transient pluripotent state during development remain elusive.

Oct4 (also known as *Pou5f1*) and Sox2 are prominent transcription factors (TFs) that regulate early embryonic development. In mice, Oct4 is expressed in all the cells of the compacted morulae, while Sox2 is first expressed in the inside cell and identified as one of the earliest markers to distinguish the inner from the outer cells (Guo *et al*., 2010; Palmieri *et al*, 1994; White *et al*, 2016). Both factors are confined to the pluripotent EPI at the late blastocyst stage (Avilion *et al*, 2003; Palmieri *et al*., 1994). *Oct4* and *Sox2* knockout (KO) leads to embryonic lethality around embryonic day (E) 4.5 and E6.0, respectively (Avilion *et al*., 2003; Nichols *et al*, 1998). Although *Oct4*- or *Sox2*-KO embryos develop apparently normal EPI, they fail to give rise to embryonic stem cells (ESCs) (Avilion *et al*., 2003; Nichols *et al*., 1998) and *Oct4*-KO embryos cannot contribute to embryonic tissues in chimera assays (Wu *et al*, 2013), suggesting that Oct4 and Sox2 are required for the EPI to acquire or maintain pluripotency. Surprisingly, *Oct4-* and *Sox2-KO* mouse embryos still express many pluripotency markers, such as Nanog, *Klf4* and *Zfp42* (*Rex1*) (Avilion *et al*., 2003; Frum *et al*, 2013; Le Bin *et al*, 2014; Stirparo *et al*, 2021; Wicklow *et al*, 2014; Wu *et al*., 2013). The controversy between the loss of pluripotency and the maintenance of the pluripotency-related genes in *Oct4-* and *Sox2-KO* EPI prompted us to investigate the role of Oct4 and Sox2 in early mouse embryos.

Recent studies suggested that Oct4 and Sox2 may function before the blastocyst stage. The long-lived presence of exogenous Oct4 and Sox2 proteins on DNA was observed in some of the 4-cell blastomeres, that tended toward the ICM rather than the TE (Plachta *et al*, 2011; White *et al*., 2016), suggesting that they may play a role in the initial lineage segregation between ICM and TE. Oct4 is also thought to contribute to the dramatic gain of chromatin accessibility at the 8-cell stage (Lu *et al*, 2016). Unlike in mouse embryos, the loss of OCT4 expression in human embryos compromises blastocyst development, leading to the downregulation of *NANOG* in the ICM and *CDX2* in the TE (Fogarty *et al*, 2017). Oct4 controls zygotic genome activation (ZGA) in zebrafish and human, but not in mice (Gao *et al*, 2018; Leichsenring *et al*, 2013). Thus, when and how Oct4 and Sox2 function in the early embryogenesis varies from species to species.

Oct4 and Sox2 play preeminent roles in maintaining the pluripotency-related transcriptional network in ESCs (Masui *et al*, 2007; Niwa *et al*, 2000). However, it remains unclear how these two TFs regulate pluripotency-related genes in embryos. The obvious obstacles are the heterogeneity and the scarcity of the preimplantation embryos. Unlike ESCs, which comprise a homogenous cell population, individual cells within embryos gradually differentiate into three cell types, EPI, PE and TE. This inherent cellular heterogeneity within embryos, coupled with the temporal variability in development across different embryos, poses a significant challenge when investigating the molecular processes using conventional molecular techniques. For instance, using the entire blastocyst or ICM for qPCR and bulk RNA-seq analysis would inevitably lead to a dilution and blur of the molecular processes within EPI (Frum *et al*., 2013; Wu *et al*., 2013). Although single-cell genomics is powerful to dissect cellular heterogeneity, it suffers from high dropouts, which may lead to inaccuracies in genomic profiles and cell identities (Minow *et al*, 2023). Due to the scarcity of embryos, it is unfeasible to compensate for the inaccuracy by increasing the number of cells. For example, single-cell RNA-seq (scRNA-seq) still faces challenges in accurately distinguishing between ICM and TE progenitors in compacted morula, as well as between EPI and PE progenitors in the early ICM (Deng *et al*., 2014; Li *et al*, 2023; Yanagida *et al*, 2022).

MEK inhibitor (MEKi) promotes the ground state of ESCs by suppressing the PE-inducing ERK signaling pathway (Ying *et al*, 2008). In embryos, MEKi has been shown to activate Nanog in ICMs, effectively suppressing the development of PE and directing the entire ICM towards a pluripotent epiblast fate (Nichols *et al*, 2009). The naïve pluripotency and readiness for differentiation of MEKi-treated EPI has been confirmed by its contribution to chimaeras with germline transmission. MEKi may thus facilitate the study of molecular events within the EPI cells by reducing the heterogeneity in ICMs.

In this study, we analyzed the transcriptome and global chromatin accessibility in the *Oct4*- or *Sox2*-KO mouse embryos using low-input RNA-seq and ATAC-seq. The effects of *Oct4* KO and *Sox2* KO are relatively small in morulae, but become evident in ICMs at the blastocyst stage. Oct4 and Sox2 mainly activate pluripotency-related genes cooperatively through the putative enhancers containing the composite OCT-SOX motifs. We observed a substantial reorganization of open chromatin regions and transcriptome in the early ICM, which was disturbed in absence of either Oct4 or Sox2. These results indicate the critical roles of Oct4 and Sox2 in establishing the pluripotency network during early mouse development.

## Results

### Generation of maternal-zygotic KO embryos

To investigate the role of Oct4 and Sox2 in the development of preimplantation embryos, we produced four transgenic mouse lines: *Oct4* KO labeled with mKO2 (monomeric kusabira-orange 2, *Oct4^mKO2^*), *Sox2* KO labeled with EGFP (*Sox2^EGFP^*), floxed *Oct4* (*Oct4^flox^*) and floxed *Sox2* (*Sox2^flox^*) (Figure 1A, 1B; Materials and methods). Although the expression of mKO2 and EGFP is driven by the constitutive mouse PGK promoter, we observed that the expression of these reporters is weak until the 8-cell and blastocyst stages, respectively, when the strong zygotic expression of Oct4 and Sox2 is observed.

**Figure 1.**
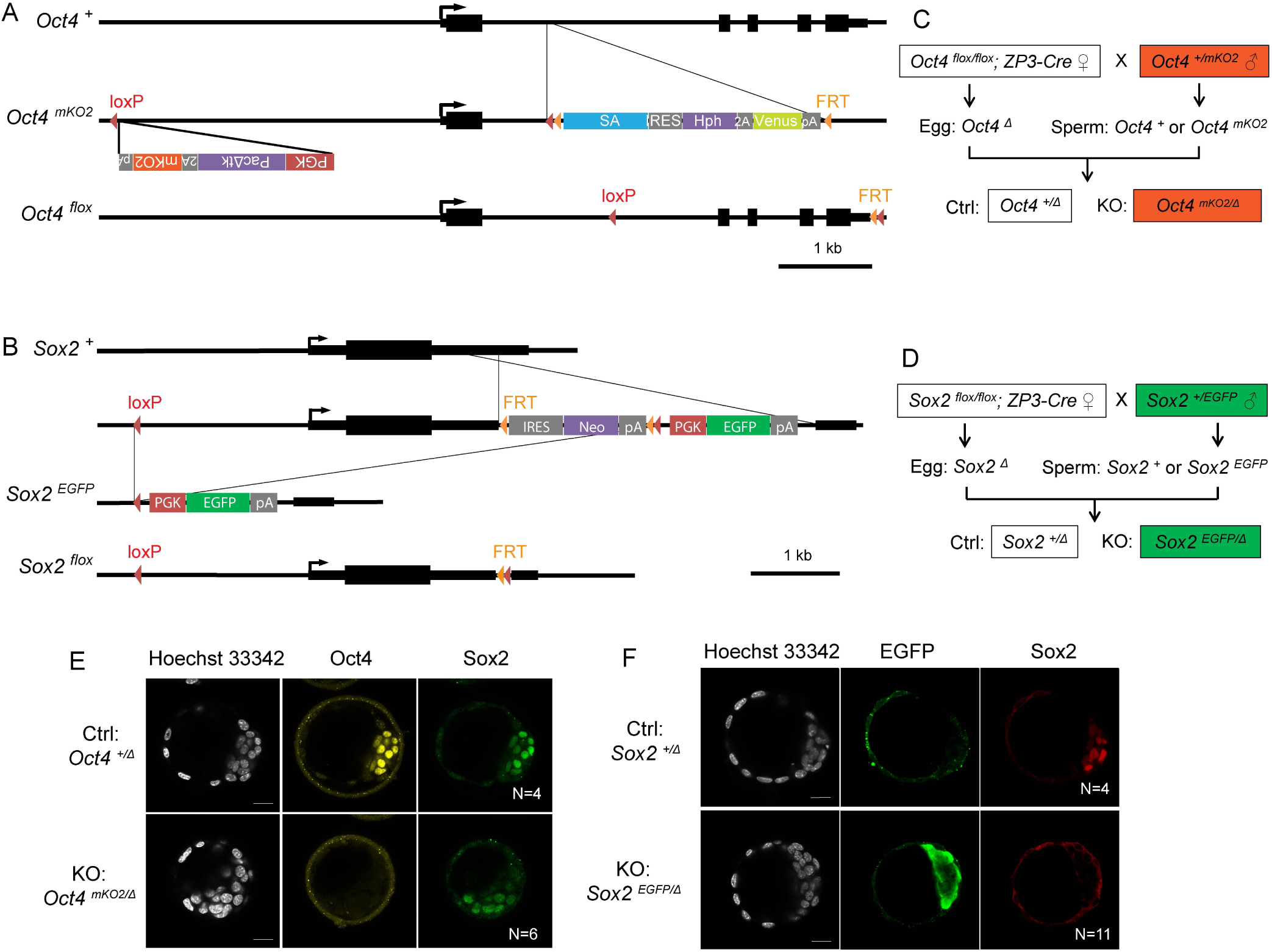
Generation of maternal-zygotic KO embryos. **A**. Schemes of mKO2-labeled *Oct4* KO (*Oct4^mKO2^*) and *Oct4 ^flox^* alleles. In the *Oct4^mKO2^* allele, a PGK-pacΔtk-P2A-mKO2-pA cassette was inserted 3.6 kb upstream of the *Oct4* transcription start site (TSS) and a promoter-less FRT-SA-IRES-hph-P2A-Venus-pA cassette was inserted into Oct4 intron 1. The inclusion of a stop codon followed by three sets of polyadenylation signal sequences (pA) after the *Venus* cassette ensures both transcriptional and translational termination, effectively blocking the expression of Oct4 exons 2–5. **B**. Schemes of EGFP-labeled *Sox2* KO (*Sox2^EGFP^*) and *Sox2 ^flox^* alleles. In the *Sox2^EGFP^* allele, the 5’ untranslated region (UTR), coding sequence and a portion of the 3’ UTR of *Sox2* were deleted and replaced with a PGK-EGFP-pA cassette. Notably, 1,023 bp of the *Sox2* 3’UTR remain intact. **C**. Mating strategy for *Oct4* Ctrl and KO embryos. **D**. Mating strategy for *Sox2* Ctrl and KO embryos. **E**. Immunostaining with embryos separated based on the mKO2 fluorescence. **F**. Immunostaining with embryos separated based on the EGFP fluorescence. N, the number of embryos. Scalebar, 20 μm. The online version of this article includes the following figure supplements for figure 1:

*Oct4 ^+/mKO2^* and *Sox2 ^+/EGFP^* heterozygous male mice were mated with *Oct4 ^flox/flox^*; *ZP3-Cre* and *Sox2 ^flox/flox^*; *ZP3-Cre* maternal KO female mice, respectively, to generate fluorescently labeled maternal-zygotic KO and unlabeled maternal KO (herein called control, Ctrl) embryos (Figure 1C, 1D). These reporter systems allow prospective identification of KO embryos using fluorescent microscopy (Figure 1-figure supplement 1), so we were able to pool embryos with the same genotype. Immunostaining results confirmed that the mKO2+ and EGFP+ embryos lost Oct4 and Sox2 proteins, respectively (Figure 1E, 1F).

### Oct4 and Sox2 regulate the chromatin landscape and transcriptome in the ICM

To assess the genome-wide molecular impact of the loss of Oct4 and Sox2, we performed low-input ATAC-seq and RNA-seq using early (E2.75) and late (E3.25) compacted morulae, as well as early (E3.75) and late (E4.5) ICMs (Materials and methods). For ATAC-seq, we pooled embryos based on our reporter systems because single-embryo samples yielded sparse signals and too few peaks (data not shown). We opted to collect single morulae or ICMs for RNA-seq, as this approach enabled us to account for embryo-to-embryo variability and detect transcripts with greater sensitivity compared to scRNA-seq (Boroviak *et al*, 2015). Given that the ICM consists of the progenitors of EPI and PE cells, we treated the embryos with the MEKi (PD0325901) from E2.5 to suppress PE development and specify the entire ICM to the EPI (Nichols *et al*., 2009) (Figure 1-figure supplement 2).

Principal component analysis (PCA) of the ATAC-seq data reveals a clear clustering of samples based on the stages and genotypes (Figure 2A). The *Oct4*-KO early and late morulae clustered close to their Ctrl counterparts, while the *Oct4-* and *Sox2-*KO early and late ICM samples clustered separately from their Ctrl. In total, 152,393 peaks were identified across different stages and genotypes. Of these, 14,016 (9.2%) and 6,637 (4.4%) showed significant decreases, while 11,884 (7.8%) and 3,660 (2.4%) exhibited significant increases in *Oct4-* and *Sox2*-KO late ICM, respectively (Figure 2B; Data supplement 1). The number of decreased peaks exceeded that of increased peaks across all the KO samples. K-means clustering of the 33,520 differential peaks identifies subsets of open chromatin regions that were differently affected by *Oct4* and *Sox2* KO (Figure 2C). Notably, compared to the peaks more reliant on Oct4 than Sox2 (Figure 2B, clusters 1 and 11), those highly reliant on both Oct4 and Sox2 (clusters 3, 8 and 14) show greater enrichment of the OCT-SOX motif. The former group tended to be already open in the morula, while the latter group became open in the ICM. Over 96.8% of the significantly changed peaks were identified as putative enhancers located distally to the transcription start sites (TSSs) (Figure 2-figure supplement 3A).

**Figure 2.**
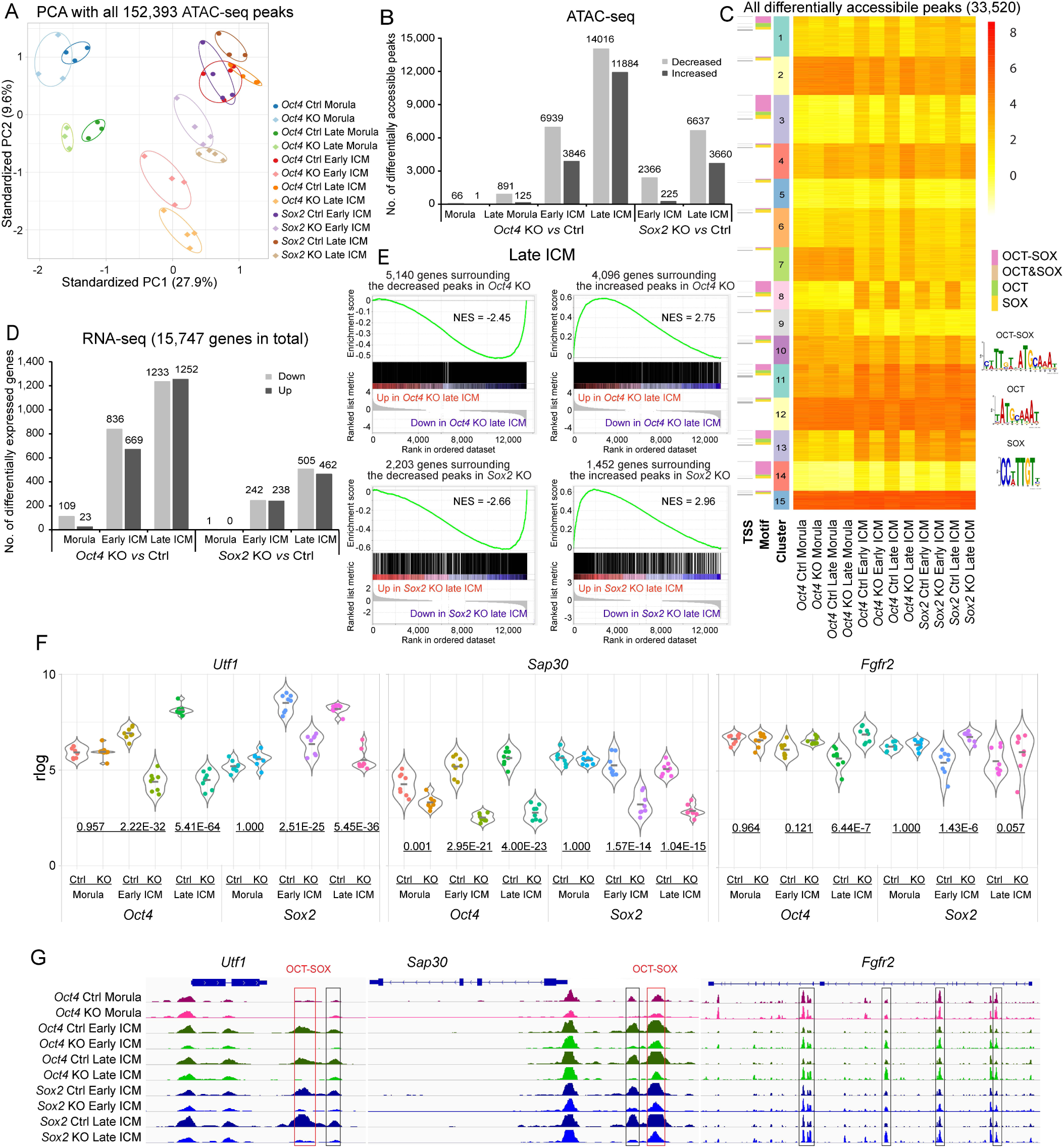
Oct4 and Sox2 regulate chromatin landscape and transcriptome in the ICM. **A**. PCA plot of all the identified ATAC-seq peaks. **B**. Number of differentially accessible ATAC-seq peaks in KO *vs* Ctrl samples. Cutoff, adjusted p-value <0.05. **C**. k-means clustering of all the significantly differential ATAC-seq peaks in KO *vs* Ctrl in Figure 2B. The heatmap is sorted by clusters, motifs and TSS. Peaks located within 100 bp from TSS were considered as TSS peaks. OCT-SOX, OCT or SOX motifs indicate that the peak contains the canonical OCT-SOX, OCT or SOX motif, respectively, while OCT&SOX motif indicates that separate OCT and SOX motifs were discovered in one peak. Cutoff, adjusted p-value <0.05. **D**. Number of differentially expressed genes in KO *vs* Ctrl samples. Cutoff, adjusted p-value <0.05 and log2 fold change ≥1. **E**. GSEA shows the correlation between significantly changed ATAC-seq peaks and the transcription of genes whose TSSs are located within 10 kb of the peak centers in *Oct4*- or *Sox2*-KO late ICMs. NES, normalized enrichment score. **F**. Examples of down- and up-regulated genes. The underlined numbers represent the adjusted p values. **G**. The ATAC-seq profiles surrounding the genes in E. Boxes mark the differentially accessible peaks, and red boxes specifically mark those with the OCT-SOX motif. The online version of this article includes the following figure supplements for figure 2:

To validate the authenticity of the Sox2-regulated downstream ATAC-seq peaks identified in this study, we compared our data to a recent study that did not employ MEKi treatment (Data ref: Li *et al*, 2023) (Li *et al*., 2023). The trends observed in our Sox2-KO late ICMs, including decreased, increased, and unchanged peaks, remained consistent in the absence of MEK inhibitor (Figure 2-figure supplement 3B). Remarkably, the decreased ATAC-seq peaks were enriched with Sox2 CUT&RUN signals (Figure 2-figure supplement 3C). Analysis of published ChIP-seq data in ESCs (Data ref: Marson *et al*, 2008; Whyte *et al*, 2012) (Marson *et al*, 2008; Whyte *et al*, 2012) shows that the peaks decreasing in the *Oct4*- or *Sox2*-KO late ICMs were frequently bound by Oct4, Sox2 and Nanog, and enriched with the active enhancer mark H3K27ac (Figure 2-figure supplement 3D). The Sox2-dependent peaks around the *Sap30* gene are shown as examples (Figure 2-figure supplement 3E). These data suggest that the decreased peaks identified in our ATAC-seq could be the potential targets of Oct4 and Sox2 in the ICM.

To assess the impact of *Oct4* and *Sox2* deletion on the transcriptome, we performed RNA-seq on individual morulae or ICMs. Mouse embryos with a maternal KO or zygotic heterozygous KO of either factor show no noticeable phenotype or molecular difference (Figure 2-figure supplement 4A) (Avilion *et al*., 2003; Frum *et al*., 2013; Kehler *et al*, 2004; Nichols *et al*., 1998; Wicklow *et al*., 2014; Wu *et al*., 2013). Therefore, we employed maternal-KO zygotic-heterozygous as our Ctrl group. In morulae, *Oct4* KO had only a minor effect on the transcriptome (Figure 2D). We observed a progressively profound impact of *Oct4* KO on the transcriptome from the morula to the late ICM, mirroring patterns at the chromatin level (Figure 2A-2D). As Sox2 is only expressed at very low levels until the blastocyst stage anyway, the transcriptome in *Sox2*-KO morulae was not significantly affected. *Oct4* or *Sox2* KO altered the expression of 2,485 (15.8%) and 967 (6.1%) genes in the late ICM, respectively (Figure 2D; Data supplement 2). *Oct4* KO had a greater effect than *Sox2* KO on both the chromatin accessibility and the transcriptome (Figure 2A-2D), suggesting Oct4 may play a more important role than Sox2. This is consistent with the earlier developmental arrest of *Oct4*-KO embryos at E4.5 compared to *Sox2*-KO embryos at E6.0. Down- and up-regulated genes identified in our *Oct4*- and *Sox2*-KO ICMs show enrichment with genes exhibiting down- and up-regulation, respectively, in the studies without MEKi (Data ref: Li *et al*., 2023; Stirparo *et al*., 2021) (Li *et al*., 2023; Stirparo *et al*., 2021) (Figure 2-figure supplement 4B-4D), confirming that MEKi did not alter the molecular functions of Oct4 and Sox2 in the ICM.

Next, we explored whether changes in chromatin accessibility affected the transcription of nearby genes, specifically those with TSSs located within 10 kb of the peak centers. Gene set enrichment analysis (GSEA) shows that the genes close to the decreased peak centers were frequently downregulated, whereas those near the increased peak centers were upregulated in the *Oct4-* and *Sox2*-KO embryos (Figure 2E; Figure 2-figure supplement 5). For example, the expression of *Utf1*, a known target of Oct4 and Sox2, as well as the accessibility of its well-characterized OCT-SOX enhancer decreased in the *Oct4-* and *Sox2*-KO ICMs (Nishimoto *et al*, 1999) (Figure 2F, 2G). On the other hand, the expression of *Fgfr2* and the accessibility of ATAC-seq peaks within its introns increased in the *Oct4*- and *Sox2*-KO ICM. In addition, integration of the ATAC-seq and RNA-seq data allowed us to infer previously unknown targets of Oct4 and Sox2, such as *Sap30* and *Uhrf1*, which are essential for somatic cell reprogramming and embryonic development (Figure 2F, 2G) (Cao *et al*, 2019; Li *et al*, 2017; Maenohara *et al*, 2017).

Taken together, the data presented so far indicate that Oct4 and Sox2 play a crucial role in shaping the chromatin landscape and transcriptome in the ICM. Consequently, we observed apparently normal development of the *Oct4*- and *Sox2*-KO embryos up to the blastocyst stage.

### Oct4 and Sox2 activate the pluripotency network in the ICM

To further explore the role of Oct4 and Sox2, we wanted to find out which genes they might influence in the ICM. As expected, many of the differential ATAC-seq peaks were consistently changed in *Oct4*- and *Sox2*-KO late ICMs (Figure 3A, Data supplement 1). However, there are a number of peaks exclusively changed in either *Oct4*- or *Sox2*-KO ICMs. The genes around the consistently decreased peaks were enriched for terms related to pluripotency and preimplantation embryonic development, such as cellular response to LIF signaling and stem cell population maintenance (Figure 3B, upper panel). The consistently elevated peaks preferentially located near the genes related to extraembryonic lineages and organogenesis, such as embryonic placenta, extraembryonic trophoblast, muscle and neural tube (Figure 3B, lower panel). Interestingly, the genes near the peaks which decreased only in the *Oct4-*KO but not in the *Sox2-*KO ICM were enriched with terms of LIF signaling and blastocyst formation (Figure 3-figure supplement 6).

**Figure 3.**
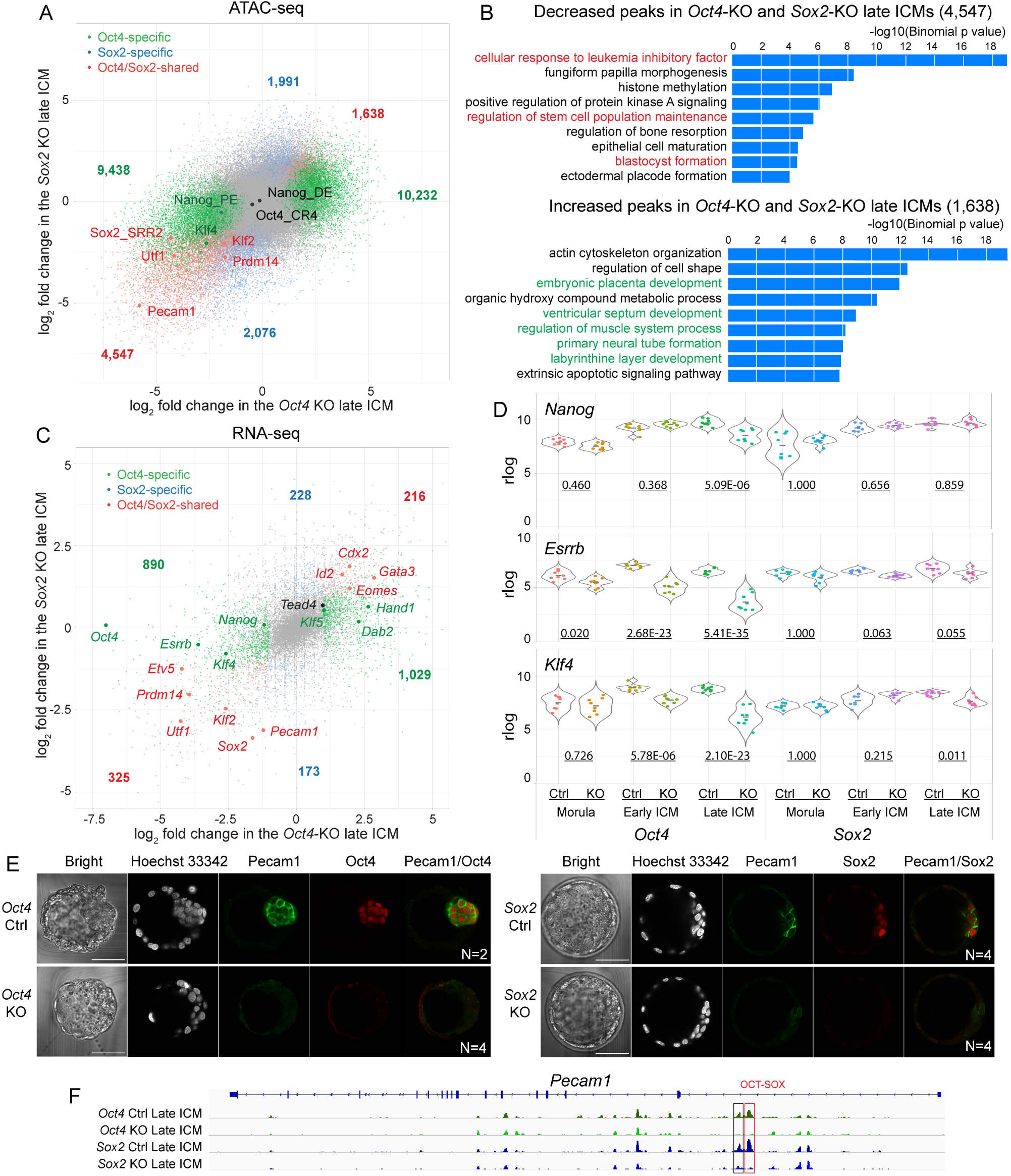
Oct4 and Sox2 activate EPI-specific genes and suppress TE-specific genes in ICM. **A**. A scatter plot shows the log_2_ fold change of ATAC-seq signals in the *Oct4*-KO and *Sox2*-KO late ICM. Colored dots represent the significantly changed peaks. Cutoff, adjusted p-value <0.05. **B**. GREAT ontology enrichment analysis of the significantly changed peaks shared in the *Oct4*-KO and *Sox2*-KO late ICM. Red terms are related to the pluripotency and preimplantation embryonic development, and green ones are related to the development of embryonic and extraembryonic lineages in the post-implantation embryos. **C**. A scatter plot shows the log_2_ fold changes of RNA-seq signals in the *Oct4*-KO and *Sox2*-KO late ICM. Cutoff, adjusted p-value <0.05 and log2 fold change ≥1. **D**. Violin plots of RNA-seq rlog values for *Nanog*, *Esrrb* and *Klf4* in the embryos. The underlined numbers represent the adjusted p values. **E**. Immunostaining of Pecam1 in the late blastocysts (E2.5+2 days). Scalebar, 50 μm. **F**. The accessibility of putative enhancers around *Pecam1* in ICMs. Boxes mark the decreased peaks and red box marks the peak with the OCT-SOX motif. The online version of this article includes the following figure supplement for figure 3:

At the transcriptional level, many known pluripotency genes, such as *Klf2*, *Etv5*, *Prdm14* and *Pecam1*, were downregulated in the *Oct4*- and *Sox2*-KO ICM (Figure 3C). In contrast, *Gata3, Cdx2* and *Eomes*, which are important for TE development and differentiation (Ralston *et al*, 2010; Russ *et al*, 2000; Strumpf *et al*, 2005), were upregulated in the *Oct4*- and Sox2-KO ICM. Interestingly, we observed that *Nanog*, *Esrrb* and *Klf4* were significantly downregulated in the *Oct4-*KO ICM, whereas they were not or only slightly downregulated in the *Sox2-*KO ICM (Figure 3C, 3D). Downregulation of *Pecam1* was confirmed at the protein level (Figure 3E). Chromatin accessibility at its putative enhancers also decreased accordingly (Figure 3F). Oct4 and Sox2 activated the components of several epigenetic modifiers, such as *Ezh2* (PRC2 H3K27me2/3 methyltransferase) and *Sap30* (a component of mSin3A histone deacetylase complex) (Figure 2F, 2G; Data supplement 2), suggesting their potential contribution to establishing the ICM-specific epigenetic status through regulation of the epigenetic modifiers.

Taken together, the above data show that Oct4 and Sox2 regulate a large number of TFs, epigenetic factors and signaling pathways in the ICM.

### Oct4 and Sox2 co-activate their targets through putative OCT-SOX enhancers

Although there is a general consensus on the cooperative binding of Oct4 and Sox2 to the OCT-SOX composite motif, the principle of the cooperation still remains controversial in different scenarios (Biddle *et al*, 2019; Chen *et al*, 2014; Friman *et al*, 2019; Li *et al*, 2019; Michael *et al*, 2020). Therefore, we investigated how Oct4 and Sox2 regulate their target open chromatin regions in the ICM. As expected, the OCT-SOX, OCT and SOX motifs were also among the most enriched motifs in the group of decreased peaks in *Oct4*- or *Sox2*-KO ICMs (Figure 4A). The motifs of other known pluripotency-related TFs, such as Klf2/4 and Esrrb, were also enriched at the decreased peaks. The elevated peaks were enriched for the GATA, TEAD, EOMES and KLF motifs, but not for the OCT-SOX, OCT or SOX motifs. Additionally, the gain of chromatin accessibility occurred at later stages compared to the loss of chromatin accessibility (Figure 2B). Thus, the increased peaks may not represent the direct targets of Oct4 or Sox2, but rather may be a secondary effect of the upregulated TE TFs, such as *Gata3*, *Tead4*, *Eomes* and *Klf5/6*, or the disruption of inhibitory interactions between Oct4/Sox2 and TE TFs (Niwa *et al*, 2005) (Figure 3C; Data supplement 2). As a control, the CTCF motif was not enriched in either the decreased or increased peaks. We confirmed the enrichment of those motifs at distal peaks and concluded that the motif analysis was not biased by the small number of the TSS peaks in the dataset (Figure 2-figure supplement 3A, Figure 4-figure supplement 7).

**Figure 4.**
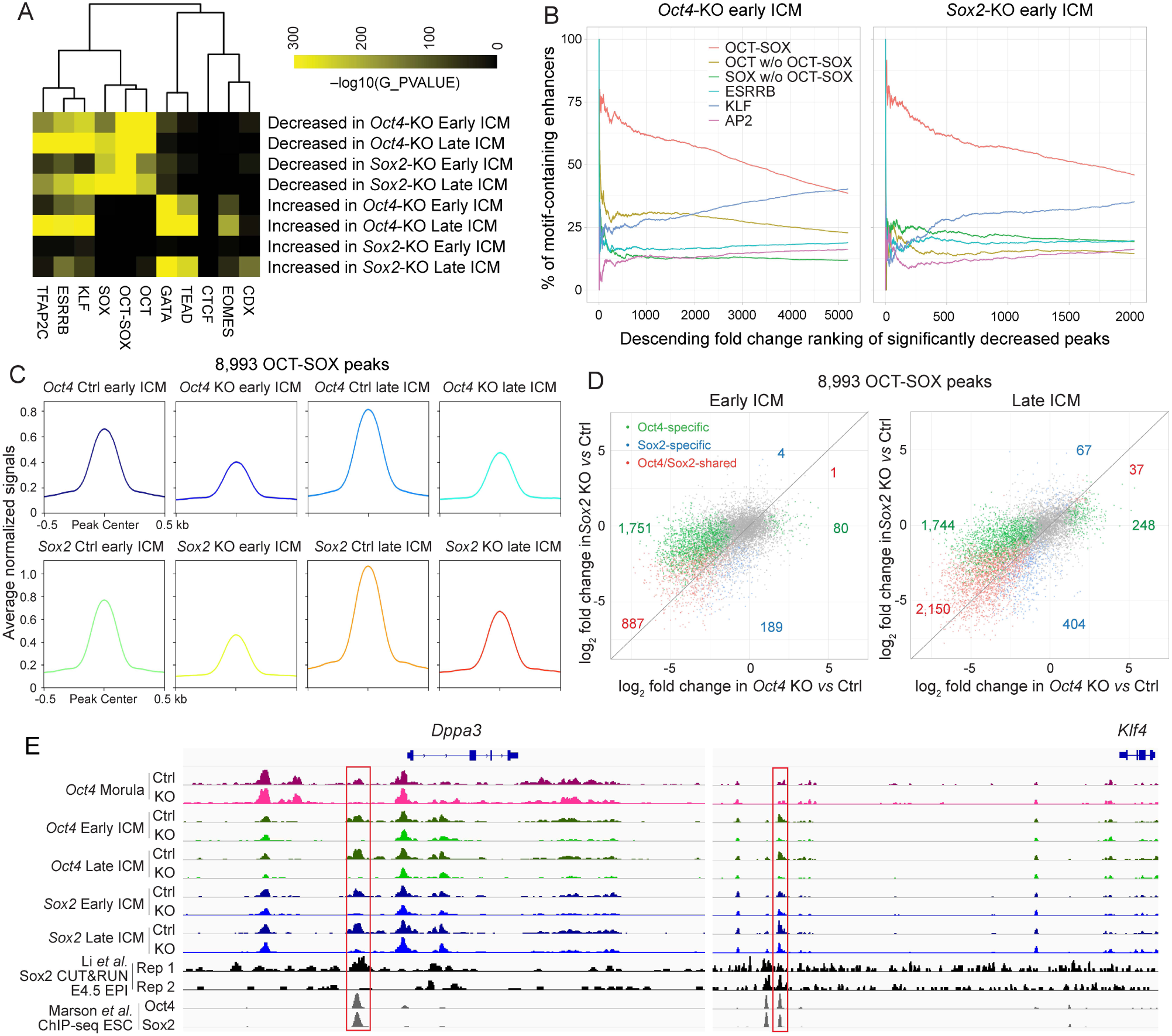
Oct4 and Sox2 activate OCT-SOX enhancers cooperatively and independently in ICM. **A**. Motif enrichment analysis of significantly changed ATAC-seq peaks in the *Oct4*-KO and *Sox2*-KO ICMs. **B**. Occurrence of the motifs in the ranked peaks. The decreased enhancers were ranked by the fold reduction. The cumulative percentages of peaks containing at least one sequence for a given motif are plotted against ranks of peaks. **C**. Average profiles of the 8,993 OCT-SOX peaks across all the Ctrl and KO ICM samples. **D**. Scatter plots show the log_2_ fold change of 8,993 OCT-SOX peaks in the *Oct4*- and *Sox2*-KO late ICMs. **E**. The profiles of ATAC-seq in early embryos, Sox2 CUT&RUN in E4.5 EPI (Li *et al*, 2023) and Oct4 and Sox2 ChIP-seq in ESCs (Marson *et al*, 2008) around the known OCT-SOX enhancers of *Klf4* and *Dppa3*. The red boxes mark the OCT-SOX enhancers. The online version of this article includes the following figure supplements for figure 4:

To investigate the effect of Oct4 and Sox2 on the chromatin accessibility, we focused on the peaks in the ICM. It is worth noting that the top decreased peaks in the *Oct4*- and *Sox2*-KO early ICM were most enriched with the OCT-SOX, but not the OCT or SOX motifs (Figure 4B), which suggests that Oct4 and Sox2 could maintain open chromatin to a greater extent at peaks containing the OCT-SOX motif (hereafter referred to as OCT-SOX peaks). The KLF motif was more enriched at the moderately decreased peaks, indicating a possible secondary effect of down-regulated *Klf2/4* (Figure 3C, 3D); alternatively, the depletion of Oct4 and Sox2 might lead to a decrease in DNA binding or the activity of Klf2/4 and other KLF TFs. In general, the accessibility of 8,993 OCT-SOX peaks decreased in both the *Oct4*- and *Sox2*-KO early and late ICM (Figure 4C; Figure 4-figure supplement 8). Furthermore, majority of the decreased OCT-SOX peaks were shared in *Oct4*- and *Sox2*-KO late ICM (Figure 4D), suggesting that both TFs participate in keeping these peaks open. The known OCT-SOX enhancers of *Dppa3* and *Klf4*, which were enriched with the binding of Oct4 and Sox2, are shown as examples (Figure 4E).

The above results suggest that the putative enhancers containing the OCT-SOX motif are primary targets of Oct4 and Sox2 in the ICM.

### Oct4 and Sox2 promote the developmental trajectory towards ICM

In early embryos, individual blastomeres are initially totipotent and indistinguishable before segregating into the ICM and TE. We subsequently investigated factors that influence the cellular trajectory toward the pluripotent ICM. Around 50% of the peaks and genes exhibiting decreased trend in *Oct4-* or *Sox2-*KO early ICMs were found to be activated in the Ctrl ICM during the natural development (Figure 5A). In alignment with earlier studies (Guo *et al*., 2010; Wu *et al*., 2016), a significant reorganization of open chromatin regions was observed upon the formation of the early ICM, resulting in the activation of 21,731 peaks (14.3%) (Figure 5B). The most prominently elevated peaks in the early ICM locate close to genes involved in cellular response to LIF, maintenance of stem cell population and blastocyst formation, in accordance with the developmental stage (Figure 5-figure supplement 9A). These peaks were enriched for OCT, SOX and OCT-SOX motifs (Figure 5-figure supplement 9B). However, when Oct4 and Sox2 were absent, the accessibility of the elevated peaks was decreased in the early ICM (Figure 5C). In alignment with the chromatin dynamics, the transcriptome also underwent substantial rearrangements in early ICMs, resulting in the upregulation of 1,115 genes (7.1%) (Figure 5D). These 1,115 genes were enriched with genes downregulated in the *Oct4-*KO and *Sox2*-KO early ICM (Figure 5E). These data indicate a critical role of Oct4 and Sox2 in initiating ICM-specific chromatin and transcriptional programs.

**Figure 5.**
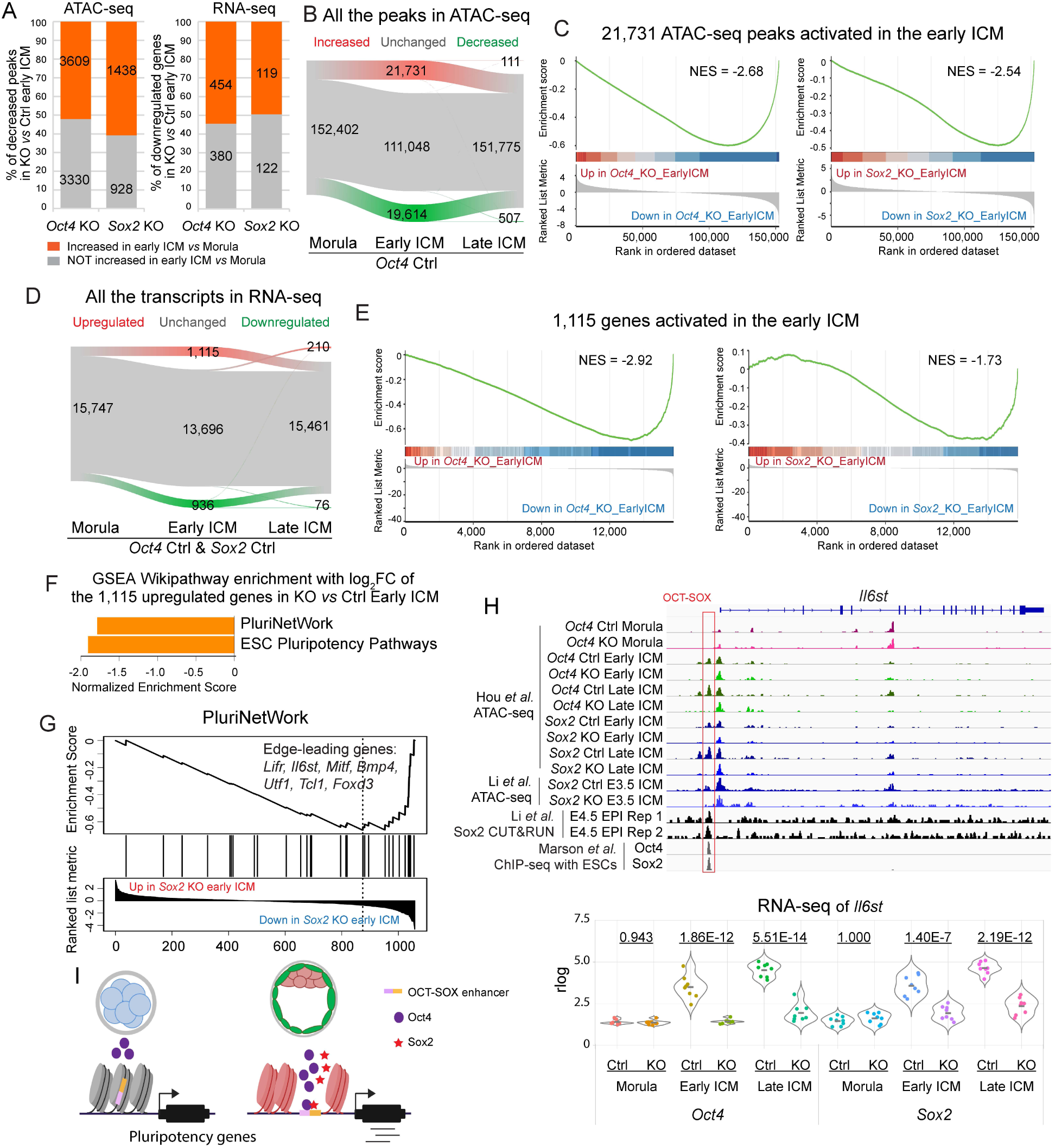
Oct4 and Sox2 promote the developmental trajectory from the morula to the ICM. **A**. Bar graphs show the dynamics of the decreased peaks (left) and genes (right) in *Oct4*- and *Sox2*-KO early ICMs from the morula to the early ICM. **B&D**. Alluvial plots show the dynamics of chromatin accessibility (B) and transcriptome (D) from morula to late ICM in the Ctrl embryos. Green, grey and red lines represent the decreased, unchanged and increased peaks/genes, respectively. In D, only genes significantly up- or down-regulated in both *Oct4* Ctrl and *Sox2* Ctrl embryos were considered as up- or down-regulated genes, while all the rest were considered as unchanged genes. **C&E**. GSEA plots show the enrichment of the 21,731 ATAC-seq peaks (C) and 1,115 genes (E) in *Oct4*-KO and *Sox2*-KO early ICMs. NES, normalized enrichment score. **F**. The bar chart illustrates the GSEA Wikipathway enrichment in WebGestalt. The log_2_ foldchange values of the 1,115 upregulated genes (Figure 5D) in *Sox2* KO *vs* Ctrl early ICM were used in this analysis. FDR≤0.05. FC, foldchange. **G**. GSEA enrichment plot of the term PluriNetWork in Figure 5F. NES, normalized enrichment score. **H**. Upper panel: IGV tracks displaying ATAC-seq and Sox2 CUT&RUN profiles (Li *et al*, 2023), along with ChIP-seq profiles of Oct4 and Sox2 in ESCs (Marson *et al*, 2008), centered around the genomic locus of *Il6st*. Red box mark the OCT-SOX enhancer. Lower panel: violin plot showing the rlog values of *Il6st*. The underlined numbers represent the adjusted p values. **I**. Model of the activation of pluripotency-related genes in the early embryos. The online version of this article includes the following figure supplements for figure 5:

The network of naïve pluripotency is governed by a core regulatory circuit of TFs, including Oct4, Sox2, Nanog, Esrrb and Klf4 (Adachi *et al*, 2018; Ng & Surani, 2011; Young, 2011). In particular, the activation of endogenous Sox2 represents a late-stage and deterministic event that triggers the fate towards pluripotency during reprogramming (Buganim *et al*, 2012). In mouse embryos, all the aforementioned TFs except Sox2 are highly expressed in the morula (Figure 5-figure supplement 9C); however, their expression does not seem sufficient to induce the mature pluripotent state at this stage. Therefore, we hypothesized that Sox2, in collaboration with Oct4, may activate the pluripotency-related genes in the early ICM. To investigate this hypothesis, we subjected the 1,115 genes and their log_2_foldchange in *Sox2*-KO and *Oct4-*KO early ICMs to GSEA Wikipathway analyses (Liao *et al*, 2019). This analysis revealed an enriched downregulation of genes associated with PluriNetWork and ESC pluripotency pathways in *Sox2*-KO early ICMs (Figure 5F). In *Oct4*-KO early ICMs, although such enrichment was not observed, a significant downregulation of pluripotency-related genes was evident (Figure 5-figure supplement 9D, 9E). For example, in the early ICM, Oct4 and Sox2 activate *Utf1* and *Il6st* (also known as *gp130*, receptor of the LIF/STAT pathway) through the open chromatin regions containing OCT-SOX motif (Figure 2F, 2G, 5H). Notably, the putative OCT-SOX enhancer of *Il6st* is enriched with Sox2 CUT&RUN signals. Additionally, Oct4 activated enzymes regulating the pyruvate metabolism (*Ldha, Me2, Pck2, etc.*) and glutathione metabolism (*Gstm1/2, Mgst2/3, Idh1*), consistent with previous studies highlighting Oct4’s role in metabolism regulation (Frum *et al*., 2013; Stirparo *et al*., 2021). These findings indicate that Sox2 expression might function as a temporal regulator in the activation of putative OCT-SOX enhancers and a subset of pluripotency-related genes in the early ICM.

In summary, above data suggest that Oct4 and Sox2 play pivotal roles in promoting the ICM fate in early mouse embryos.

## Discussion

Oct4 and Sox2 are key TFs for preimplantation embryonic development across species (Daigneault *et al*, 2018; Fogarty *et al*., 2017; Gao *et al*, 2022; Lee *et al*, 2013; Leichsenring *et al*., 2013). However, because of the paucity of embryonic cells, the timing and mechanisms of their function in regulating the EPI lineage remain unclear. In this study, we explored the roles of Oct4 and Sox2 in mouse preimplantation embryos using transgenic models.

Previous reports suggested that Oct4 and Sox2 could direct 4-cell blastomeres towards ICM fate (Plachta *et al*., 2011; White *et al*., 2016), and Oct4 facilitates chromatin opening at the 8-cell stage (Lu *et al*., 2016). Unexpectedly, our data showed that either *Oct4-*KO or *Sox2*-KO have minimal impact on global chromatin accessibility and transcription until the early blastocyst stage (Figure 2A-2D). Oct4 is expressed at similar levels between morulae and ICMs; in contrast, Sox2 mRNA and protein levels are negligible in morulae, but are significantly upregulated in ICMs (Figure 5-figure supplement 9C) (Palmieri *et al*., 1994; Wicklow *et al*., 2014). Therefore, in morulae, the function of Oct4 is likely limited by the insufficient levels of Sox2 and possibly other necessary factors. Another plausible limiting aspect could be epigenetic constraints. The epigenetic state and 3-dimensional structure of the chromatin in the morula differ from those in the ICM, which may hinder Oct4 function (Du *et al*, 2017; Wang *et al*., 2018; Wang *et al*, 2014; Zhang *et al*., 2016).

Endogenous Oct4 is activated prior to Sox2 in both embryogenesis and reprogramming. In mouse embryos, Oct4 is initially highly expressed in all blastomeres from the 8-cell stage and later restricted to the EPI, whereas Sox2 is repressed by the Hippo pathway in early stages and is specifically expressed in the EPI (Frum *et al*, 2019; Guo *et al*., 2010; Wicklow *et al*., 2014). In reprogramming, the activation of endogenous Sox2 occurs later than the endogenous Oct4 and signifies the final phase of iPSC generation (Buganim *et al*., 2012). In both scenarios, the expression of Sox2 coincides with the setup of the pluripotent state. Our data show that *Sox2*-KO ICMs fail to fully activate the pluripotent program (Figure 3C). Recently, we reported that the loss of Sox2 might contribute to decreased developmental potential of pluripotent cells upon priming (MacCarthy *et al*, 2024). Above studies suggest that Sox2 plays a critical role in regulating the spatial-temporal full onset of pluripotency-related genes.

Loss of Oct4 and Sox2 impairs embryonic pluripotency, as evidenced by the inability of KO embryos to give rise to ESCs or contribute to the embryonic part in chimera assays (Avilion *et al*., 2003; Nichols *et al*., 1998; Wu *et al*., 2013). To date, only a few of pluripotency-related genes have been found downstream of Oct4 and Sox2 in mouse embryos. Here, we found that a large number of pluripotency-related genes were downregulated in *Oct4*-KO ICM, contrasting with previous reports which suggested that most pluripotency-related genes were maintained (Frum *et al*., 2013; Le Bin *et al*., 2014; Stirparo *et al*., 2021; Wu *et al*., 2013). This discrepancy may be due to differences in the experimental design. First, unlike previous study that used the whole blastocysts (Frum *et al*., 2013), we isolated an EPI cell population through immunosurgery and short-term inhibition of MEK while it did not alter Oct4/Sox2 target open chromatin regions and genes (Nichols *et al*., 2009) (Figure 1-figure supplement 2; Figure 2-figure supplement 3B-3D, 4B-4D). Second, we removed *Oct4* exons 2 to 5 (the whole DNA-binding POU domain), resulting in a complete ablation of Oct4 function (Figure 1A). In contrast, majority of the previous studies removed only exon 1 (Frum *et al*., 2013; Wu *et al*., 2013), which encodes for N-terminal transactivation domain. The remaining exons 2 to 5 might be translated and could retain partial function. Finally, unlike the previous study that used maternal-WT zygotic-KO embryos (Stirparo *et al*., 2021), we used maternal-zygotic-KO embryos (Figure 1C). The residual levels of maternal Oct4 could have potentially rescued the KO.

Preimplantation embryos undergo dynamic epigenomic reprogramming, governed by multiple epigenetic modifiers. For example, *Ezh2* is required for the establishment and maintaining global H3K27me3, while *Uhrf1* maternal-KO embryos fail to form healthy morula due to disrupted DNA methylation and histone modifications (Cao *et al*., 2019; Maenohara *et al*., 2017; Puschendorf *et al*, 2008). Our data show that Oct4 and Sox2 activate components of epigenetic modifiers like *Ezh2*, *Uhrf1* and *Sap30* (Figure 2F, 2G; Figure 2-figure supplement 3E; Data supplement 2), suggesting their role in regulating epigenetic status.

While the cooperation of Oct4 and Sox2 on the OCT-SOX enhancers is widely acknowledged, there are still debates regarding their precise roles (Biddle *et al*., 2019; Chen *et al*, 2008; Han *et al*, 2022; Li *et al*., 2019; Michael *et al*., 2020; Velychko *et al*, 2019). In this study, most OCT-SOX peaks are more affected in *Oct4-*KO than in *Sox2*-KO ICMs, indicating that Oct4 plays a more important role than Sox2 in activating putative OCT-SOX enhancers *in vivo*. This observation aligns with a previous study showing that the forced expression of Oct4 can rescue pluripotency in *Sox2*-null ESCs (Masui *et al*., 2007). We believe that this outcome is unlikely to be attributed to compensation from other members in the Sox family. Sox1, Sox3, Sox15 and Sox18 are promising candidates as they can either rescue the *Sox2*-KO mESCs or replace Sox2 in reprogramming (Nakagawa *et al*, 2008; Niwa *et al*, 2016). However, our analysis reveals that *Sox1*, *Sox3* and *Sox18* are expressed at extremely low levels in wildtype and Sox2-KO embryos (Data supplement 2), suggesting that they are unlikely to be able to fulfill Sox2’s role (Deng *et al*., 2014). Sox15 exhibits indistinguishable expression levels in all three cell types at the blastocyst stage. Furthermore, *Sox15* KO mice displays normal health and fertility (Lee *et al*, 2004). Nonetheless, we cannot completely rule out the possibility of compensation by Sox15 in *Sox2*-KO embryos.

Our results highlight the crucial role of Oct4 and Sox2 in establishing the transcriptome and chromatin state in the pluripotent EPI. At the blastocyst stage, Oct4 and Sox2 work together to open the putative OCT-SOX enhancers and activate the pluripotency-related genes (Figure 5I). The absence of Sox2 and other factors likely limits the function of Oct4 in morulae. However, the upstream factors driving the activation of the core pluripotency regulatory circuitry remain unknown. Further studies are needed to deepen our understanding of the molecular mechanisms governing the embryonic pluripotency program and the early lineage segregation.

## Materials and methods

### Mice

To generate ESCs carrying a floxed *Oct4* allele (*Oct4^flox^*), the construct containing floxed *Oct4* exon 2-5 and a promoter-less FRT-IRES-βgeo-pA cassette was electroporated into germline-competent Acr-EGFP ESCs. Clones were screened for homologous recombination and transiently transfected with an FLP expression vector to remove the FRT cassette. To generate ESCs carrying a mutant *Oct4* allele linked to *mKO2* (*Oct4^mKO2^*), Acr-EGFP ESCs were targeted with the construct containing a PGK-pacΔtk-P2A-mKO2-pA cassette 3.6 kb upstream of the *Oct4* TSS and a promoter-less FRT-SA-IRES-hph-P2A-Venus-pA cassette in *Oct4* intron 1. To generate ESCs carrying the floxed *Sox2* allele (*Sox2^flox^*), the construct containing the 5’ loxP site 1.9 kb upstream of *Sox2* TSS and a promoter-less FRT-IRES-Neo-pA cassette followed by the 3’ loxP site in the 3’ UTR of *Sox2* was electroporated into Acr-EGFP ESCs. Clones were screened for homologous recombination and transiently transfected with an FLP expression vector to remove the FRT cassette. To generate ESCs carrying a conditional allele of *Sox2* linked to *EGFP* (*Sox2^EGFP^*), Acr-EGFP ESCs were targeted with the construct containing the 5’ loxP site 1.9 kb upstream of *Sox2* TSS, and a promoter-less FRT-IRES-Neo-pA cassette, the 3’ loxP site, and a PGK-EGFP-pA cassette in the 3’ UTR of *Sox2*. Chimeric mice were generated by morula aggregation and heterozygous mice were obtained through germline transmission. The Zp3-Cre transgenic mice were used to generate maternal KO alleles. Animal experiments and husbandry were performed according to the German Animal Welfare guidelines and approved by the Landesamt für Natur, Umwelt und Verbraucherschutz Nordrhein-Westfalen (State Agency for Nature, Environment and Consumer Protection of North RhineWestphalia). The work was funded by the Max Planck Society and the Max Planck Society’s White Paper-Project “Animal testing in the Max-Planck-Society”.

### Embryo collection

All the embryos for RNA-seq and ATAC-seq were collected from female mice superovulated with PMSG and hCG. Early and late morula samples were collected at E2.75 and E3.25, respectively. Early and late ICM samples were collected at E2.5 and cultured in KSOM with MEKi (PD0325901, 1 μM) until early and late blastocyst stages, respectively.

### ICM isolation

Zona pellucida was removed by brief incubation in prewarmed acidic Tyrode’s solution. ICM was obtained by immunosurgery. Briefly, blastocysts were incubated in 20% rabbit anti-mouse whole serum (Sigma-Aldrich) in KSOM at 37°C, 5% CO2 for 15 minutes, followed by three rinses in M2. Afterwards, embryos were incubated in 20% guinea pig complement serum (Sigma-Aldrich) in KSOM at 37°C, 5% CO2 for 15 minutes, followed by three rinses in M2. In the end, ICM was isolated by repetitive blowing in mouth pipette to remove the debris of dead trophectoderm cells.

### Immunofluorescence

Immunofluorescence was performed using the following primary antibodies with dilutions: mouse monoclonal anti-Oct4 (sc-5279, Santa Cruz), 1:1000; goat anti-Sox2 (GT15098, Neuromics), 1:300; rabbit anti-mKO2 (PM051, MBL), 1:1000; rabbit anti-GFP (ab290, Abcam), 1:500; goat anti-Pecam1 (AF3628, R&D biosystems), 1:300; rat monoclonal anti-Nanog (14-5761-80, Thermo Fisher Scientific), 1:100; goat anti-Sox17 (AF1924, R&D Systems), 1:1000 of 0.2 mg/ml.

### ATAC-seq and data analysis

ATAC-seq libraries were prepared as previously described with some modifications. 20 to 40 morulae or ICMs were pooled and lysed in the lysis buffer (10 mM Tris·HCl, pH=7.4; 10 mM NaCl; 3 mM MgCl2; 0.15% Igepal CA-630; 0.1% Tween-20) for 10 min on ice. We added 0.1% Tween-20 to the lysis buffer as it greatly reduced reads mapped to mitochondrial DNA. Lyzed embryos were briefly washed, collected in 2.0 μl PBS containing 0.1 mg/ml polyvinyl alcohol (PVA) and transferred to 3.0 μl Tn5 transposome mixture (0.25 μl Tn5 transposome, 2.5 μl tagmentation buffer, Illumina, FC-121-1030; 0.25 μl H_2_O). The samples were incubated in 37 °C water bath for 30 min. Tagmentation was stopped by adding 2.0 μl 175 mM EDTA and incubated at 50 °C for 30 min. Excess EDTA was quenched by 2.0 μl 160 mM MgCl_2_. The libraries were amplified in the following reaction: 9.0 μl Transposed DNA, 10.0 μl NEBNext High-Fidelity 2×PCR Master Mix (New England Biolabs, M0541S), 0.25 μl 100 μM PCR Index 1, 0.25 μl 100 μM PCR Index 2 and 0.5 μl H_2_O. The sequences of index 1 and 2 are in Table supplement 2. The PCR program is as the following: 72 °C for 5 min; 98 °C for 30s; 16 thermocycles at 98 °C for 10 s, 63 °C for 30 s and 72 °C for 1 min; followed by 72 °C 5 min. Amplified libraries were purified twice with 1.2 × AMPure XP beads. The sequencing was performed on the NextSeq 500 system with pair-end 75bp.

Sequence reads were trimmed for adapter sequences using SeqPurge and trimmed reads no shorter than 20 bases were aligned to the mm10 mouse reference genome using Bowtie2 with a maximum fragment size of 2000. Duplicated reads were removed using Picard MarkDuplicates (https://broadinstitute.github.io/picard/) and only reads uniquely mapped to the standard chromosomes except chrY and chrM (mitochondrial genome) with mapping quality of at least 30 were used for the following analysis. Reads on the positive and negative strands were shifted +4 bp and −5 bp, respectively. Peak calling was performed using MACS2 with the following parameters: --keep-dup all -g mm --nomodel --shift −50 --extsize 100 -B -- SPMR. Peaks that overlap with the blacklisted regions (https://sites.google.com/site/anshulkundaje/projects/blacklists; https://sites.google.com/site/atacseqpublic/atac-seq-analysis-methods/mitochondrialblacklists-1) and satellite repeats (RepeatMasker) were removed. Significant peak summits (q-value ≤0.001) from biological replicates were merged for each sample group using BEDTools with a maximum distance between two summits to be merged of 200 bp. Only merged peak summits that overlap with summits from at least two replicates were retained. Reads per million (RPM)-normalized pileup signals in the bedGraph format were converted into bigWig files using the UCSC bedGraphToBigWig tool. The bigWig files for the average signal of biological replicates were generated using the UCSC bigWigMerge tool. The heatmaps of RPM-normalized pileup signals around the TSSs of GENCODE (vM23) protein-coding genes were generated using deepTools.

For count-based analysis of transposon insertion events, merged peak regions were generated by combining the adjacent peak summits of all embryo groups and selecting 200-bp regions around the centers of each merged summit. 5’ ends of shifted read alignment on both the positive and negative strands were considered as transposon insertion sites. The number of transposon insertions in each merged peak region were counted for each sample using BEDTools and normalized and transformed to log_2_ scale using the rlog function of the Bioconductor package DESeq2 and significantly differential peaks were identified with an adjusted p-value cutoff of 0.05. Peaks located within 100 bp from TSSs of GENCODE (vM23) genes were considered as TSS-proximal peaks. The presence of TF binding motifs in merged peaks was investigated using FIMO with the default cutoff (p-value <10^-4^).

The following JASPAR motifs were used: MA0036.3 GATA2, MA0139.1 CTCF, MA0141.3 ESRRB, MA0142.1 Pou5f1-Sox2, MA0143.3 Sox2, MA0524.2 TFAP2C, MA0599.1 KLF5, MA0792.1 POU5F1B, MA0800.1 EOMES, MA0808.1 TEAD3 and MA0878.1 CDX1.

Significantly enriched motifs in each peak cluster were identified from using PscanChIP (Zambelli *et al*, 2013). Functional annotation of gene sets associated with ATAC-seq peaks was performed using GREAT (McLean *et al*, 2010) with default settings.

### RNA-seq and data analysis

Single-embryo RNA-seq was performed as previously described with some modifications (Kurimoto et al. 2007, Nakamura et al. 2015). 2,493 copies of ERCC RNA Spike-In (Ambion) were added to each single-embryo sample. To remove PCR duplicates, we used R2SP-UMI-d(T)24 primer, instead of V1d(T)24 primer, for reverse transcription (RT). To reduce byproducts derived from the RT primer, the poly(A) tailing reaction was performed only for 1 min at 37°C. cDNA was amplified with V3d(T)24 and R2SP primers and Terra PCR Direct Polymerase (Clontech) for 16 cycles. The cDNA was further amplified for four cycles with NH2-V3d(T)24 and NH2-R2SP primers, and purified thrice with 0.6x AMPure XP beads. The 3’ end enriched libraries were constructed using KAPA HyperPlus Library Preparation Kit (VWR International, KK8513) with 6 cycles of PCR. All the oligos used in the RT and library amplification are in Table supplement 3. The sequencing was performed on the NextSeq 500 system with single-end 75bp.

UMI sequence of each read was extracted from FASTQ files for Read 2 using UMI-tools. Poly-A sequences were trimmed from the 3’ end of Read 1 using Cutadapt (https://cutadapt.readthedocs.org/). The trimmed Read 1 sequences were aligned to the mm10 mouse reference genome using TopHat2 with GENCODE (vM23) transcripts as a transcriptome reference and with default parameters. The reads were also mapped to ERCC reference sequences using Bowtie2 with default parameters. To confirm the genotype of *Oct4*- and *Sox2*-KO embryos, we also mapped the reads to the PGK-pacΔtk-P2A-mKO2-hGHpA or PGK-EGFP-hGHpA cassette. The reads mapped to the genome were assigned to GENCODE transcripts using featureCounts. The number of unique transcripts assigned to each GENCODE gene was calculated using TRUmiCount. The transcript counts were normalized and transformed to log_2_ scale using the rlog function of the Bioconductor DESeq2 package. Differential expression analysis was performed on genes that were detected in at least half of the samples in at least one stage using DESeq2 and differentially expressed genes were identified with an adjusted p-value <0.05 and log_2_ fold change ≥1.

### GSEA

Gene set enrichment analysis (GSEA) (Subramanian *et al*, 2005) was performed to determine whether genes close to the selected ATAC-seq peaks are enriched in genes differentially expressed in *Oct4*- or *Sox2-*KO embryos. The ATAC-seq peaks were annotated to Ensembl protein-coding genes whose TSSs are located within 10 kb of the peak centers using the R package ChIPpeakAnno (Zhu *et al*, 2010). False discovery rate (FDR) was estimated by gene set permutation tests. Heatmaps were generated from normalized enrichment scores (NESs) for gene sets with FDR ≤0.1.

## Supporting information

Table supplement 1. Coordinates for the marked well-known enhancers in Figure 3A

Table supplement 2. oligos for ATAC-seq library preparation

Table supplement 3. oligos for RNA-seq library preparation

Dataset supplement 1. DESeq2 of ATAC-seq peaks

Dataset supplement 2. DESeq2 of RNA-seq

## Acknowledgments

We appreciate the invaluable assistance from Ingrid Gelker, Claudia Ortmeier, David Obridge, Martina Sinn and the animal facility. Special thanks go to Dong Han, Rui Fan, Eva Kutejova and Erik Tolen for their insightful discussions. We acknowledge the generous support from Anika Witten, Christoph Bartenhagen and Carolin Walter of the Core Facility Genomics at the University of Muenster for their support with sequencing. Additionally, we would like to express our gratitude to Shixue Gou and Hui Zhang from the Guangzhou National Laboratory for their kind suggestions on RNA-seq data analysis and manuscript revisions, respectively. This work was supported by the Max Planck Society.

## Additional information

### Data Availability

ATAC-seq and RNA-seq data have been deposited at GEO under GSE264614 and GSE264615, respectively. Published high-throughput sequencing datasets used in this manuscript are listed as follows: ATAC-seq and CUT&RUN of early embryos, GSE203194; scRNA-seq of early embryos, GSE203194 and GSE159030; Oct4/Sox2/Nanog ChIP-seq of mESCs, GSE11724; H3K27ac ChIP-seq of mESCs, GSE27844.

### Author contributions

K.A. and Y.H. designed the study. Y.H. performed the experiments, assembled and interpreted data, and wrote the manuscript. K.A. performed the bioinformatic analysis, interpreted the data and edited the manuscript. G.W. generated the transgenic mice and advised on the project. Z.N. and S.H. assisted with the experiments. Y.H. and Q.J. contributed to the bioinformatic analysis. S.V. and I.B. advised on the project and edited the manuscript. H.R.S. supervised the project and edited the manuscript.

## Supplementary files

**Table supplement 1.** Coordinates for the marked well-known enhancers in Figure 3A

**Table supplement 2.** oligos for ATAC-seq library preparation

**Table supplement 3.** oligos for RNA-seq library preparation

**Dataset supplement 1.** DESeq2 of ATAC-seq peaks

**Dataset supplement 2.** DESeq2 of RNA-seq

---

**Figure supplement 1.**
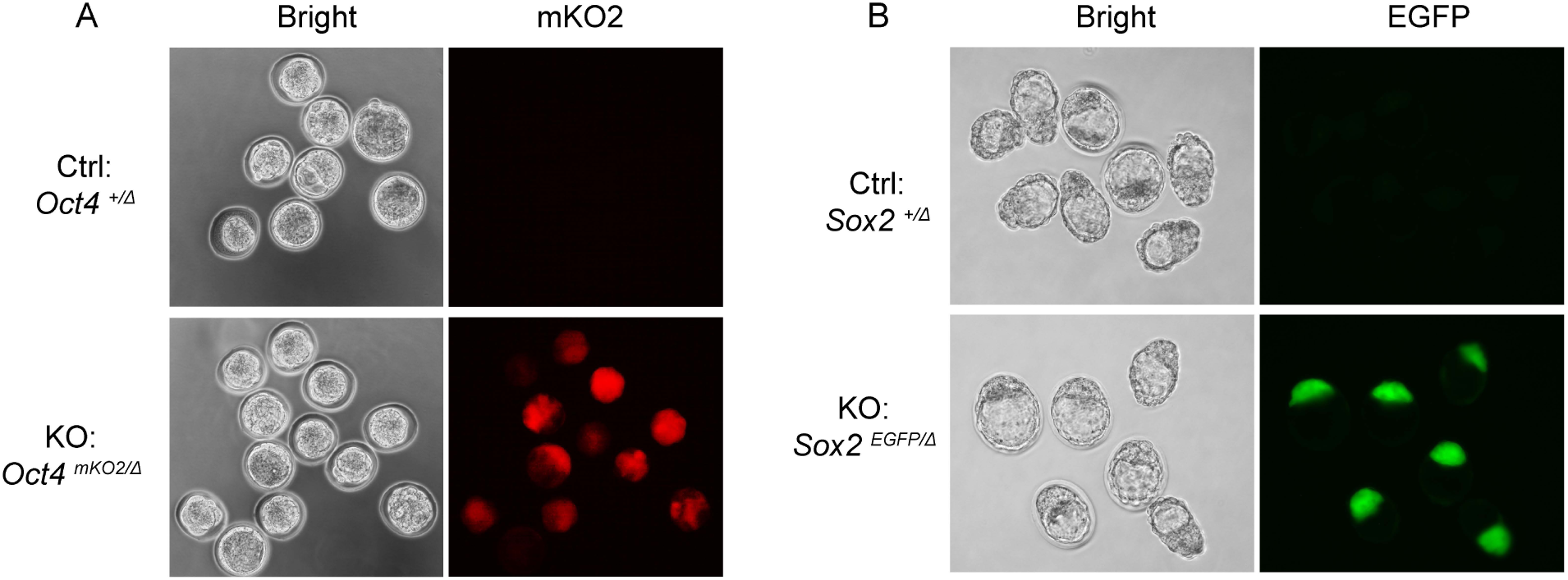
Validation of the transgenic embryos (related to Figure 1) **A**. The mKO2-labeled *Oct4*-KO embryos were identified under fluorescent microscopy at E2.5+1 day. **B**. The EGFP-labeled *Sox2*-KO embryos were identified under florescent microscopy at E2.5+2 days.

**Figure supplement 2.**
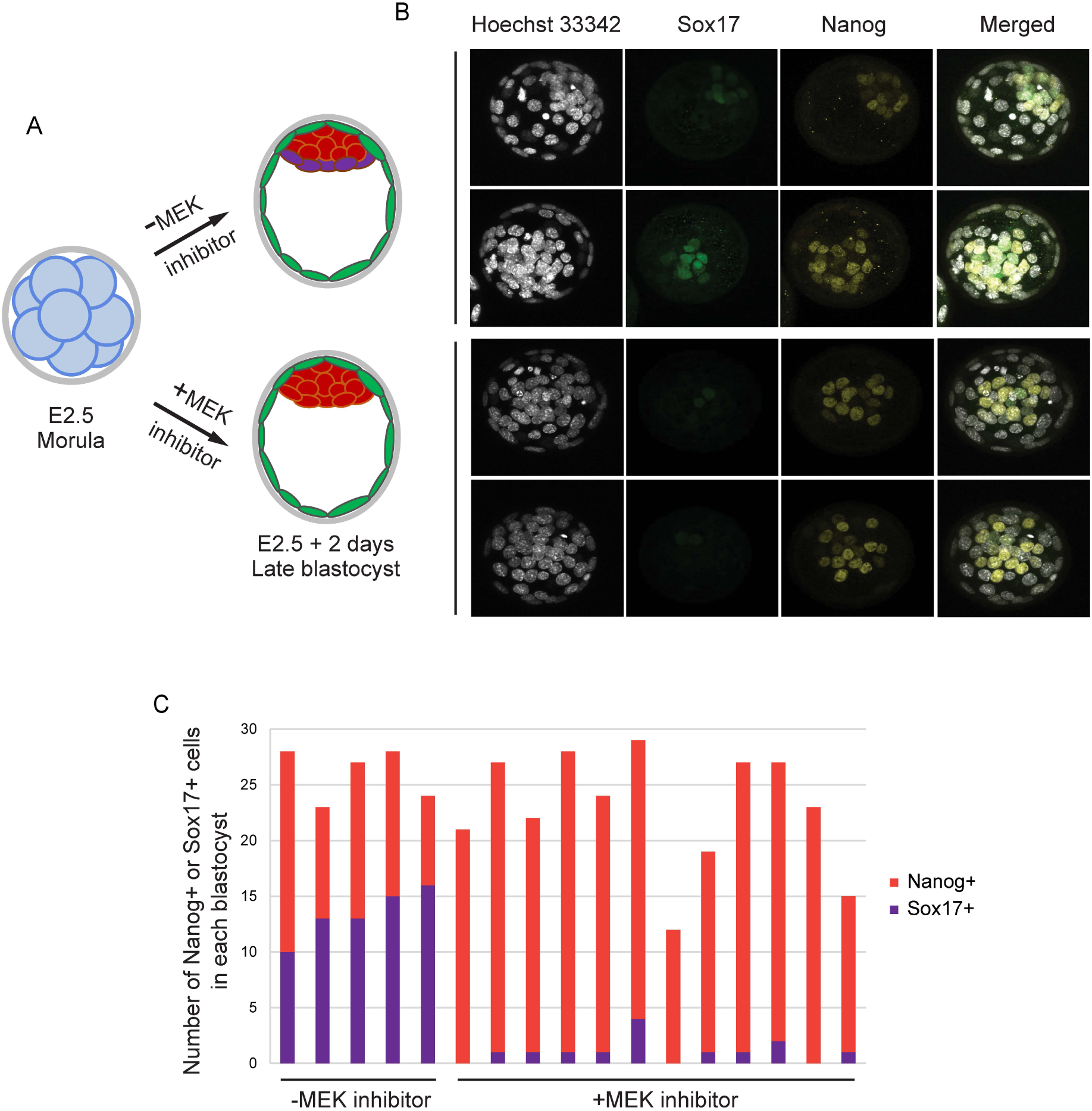
Validation of the effect of MEK/ERK inhibitor (MEK/ERKi) (related to Figure 1) **A**. The scheme of MEK/ERKi (PD0325901, 1 μM) treatment. **B**. Immunostaining of Sox17 and Nanog in the late blastocysts treated with or without MEK/ERKi. **C**. The number of Sox17+ or Nanog+ cells in each blastocyst.

**Figure supplement 3.**
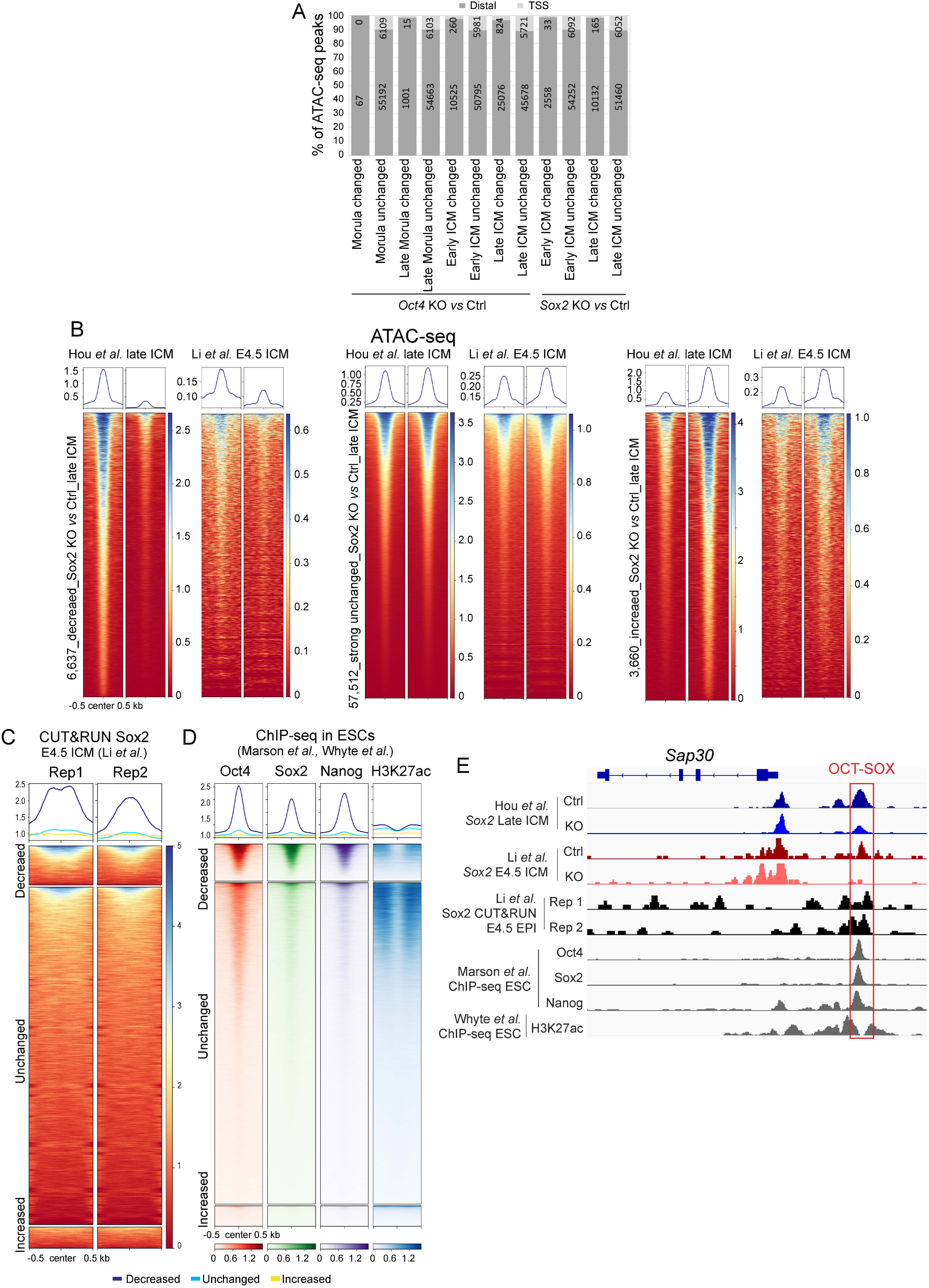
Quality check of the low-input ATAC-seq (related to Figure 2) **A**. The location of ATAC-seq peaks relative to TSSs. The number in each bar represents the number of peaks in each group. **B&C.** ATAC-seq (B) and Sox2 CUT&RUN (C) profiles (Li et al, 2023) over the decreased, unchanged and increased ATAC-seq peaks identified in our *Sox2-*KO late ICMs. **D**. ChIP-seq profiles of Oct4, Sox2, Nanog and H3K27ac in ESCs (Marson et al, 2008) over the decreased, unchanged and increased ATAC-seq peaks in our *Sox2*-KO late ICMs. To exclude spurious peaks, only strong unchanged peaks (57,512 out of 142,096) were used in the analysis from C to E. **E**. IGV tracks displaying ATAC-seq and Sox2 CUT&RUN profiles (Li *et al*, 2023) in late ICMs, along with ChIP-seq profiles of Oct4, Sox2, Nanog and H3K27ac in ESCs (Marson et al, 2008; Whyte et al, 2012), centered around the genomic locus of *Sap30*. Arrows mark the decreased ATAC-seq peaks.

**Figure supplement 4.**
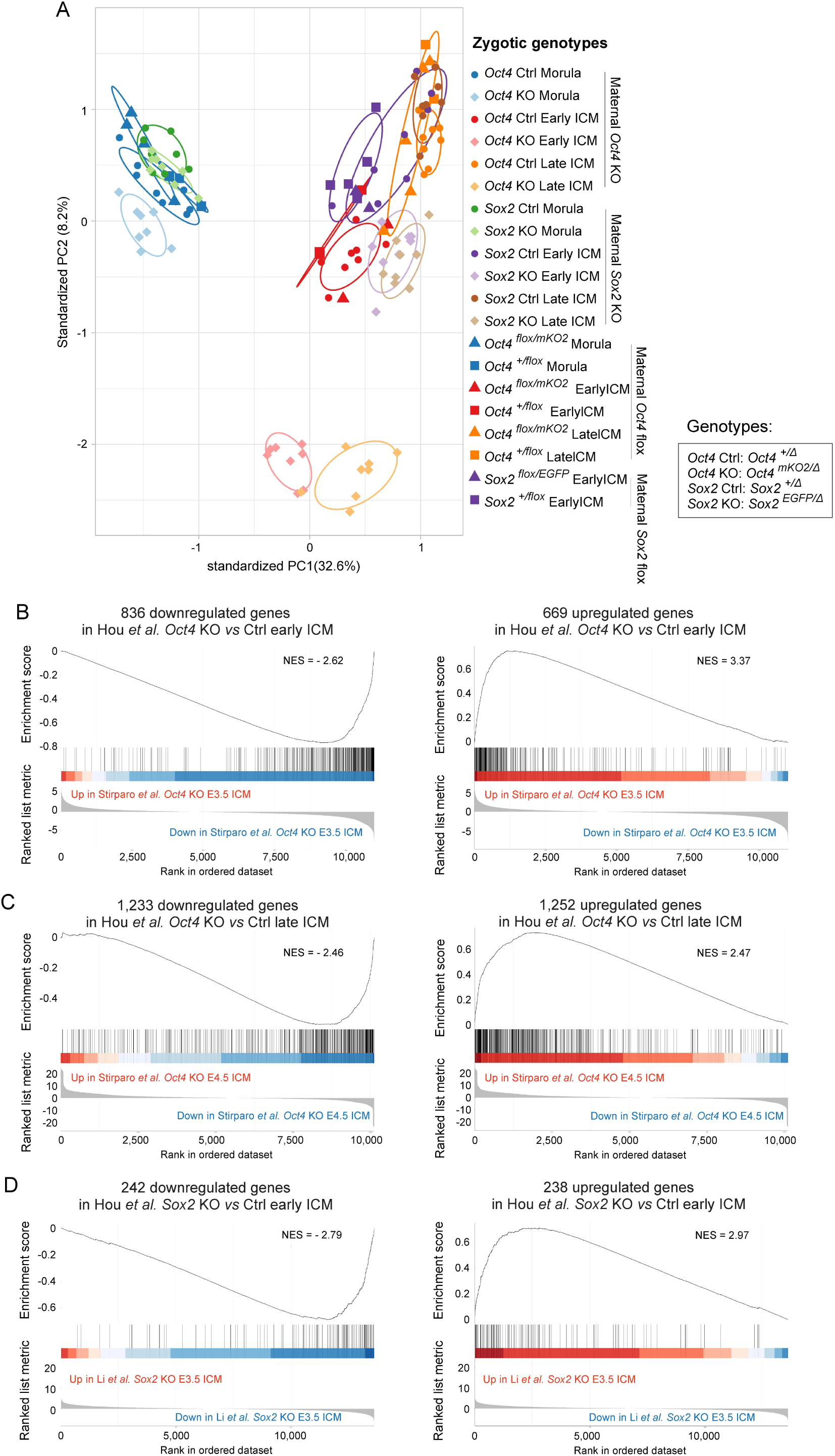
Quality check of the low-input RNA-seq (related to Figure 2) **A**. PCA plot illustrating the distribution of RNA-seq samples with the indicated stages and genotypes. **B-D**. GSEA plots of down- and up-regulated genes identified in our *Oct4*-KO early ICM (B), *Oct4*-KO late ICM (C) and *Sox2*-KO early ICM (D) in the scRNA-seq dataset of ICM samples without MEK/ERKi (Stirparo *et al*., 2021; Li *et al*., 2023). NES, normalized enrichment score.

**Figure supplement 5.**
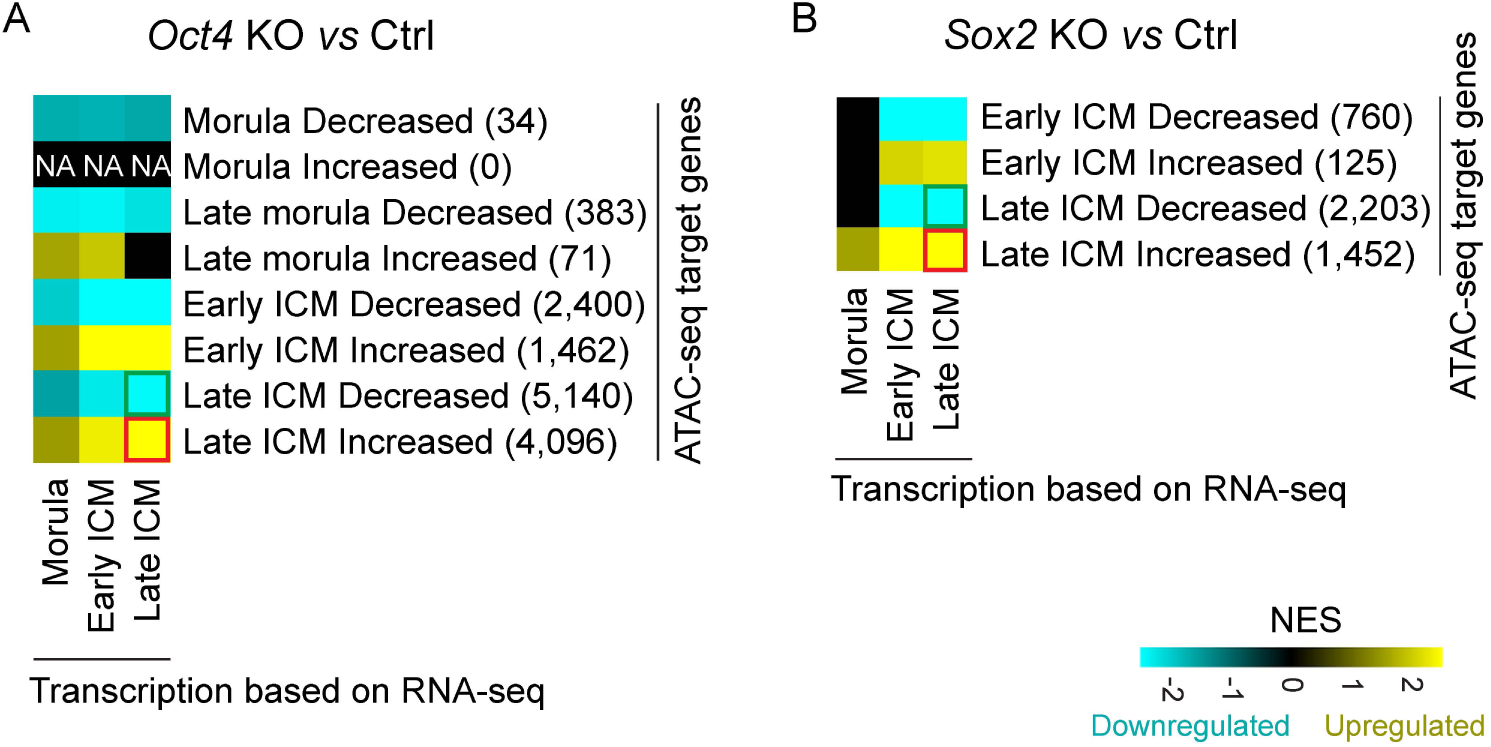
The correlation between chromatin accessibility and the transcription activity of surrounding genes (related to Figure 2) Heatmap of GSEA enrichment *Oct4*-KO (A) and *Sox2-*KO (B) samples. The analysis utilized genes with TSS falling within a 10 kb proximity to the decreased or increased ATAC-seq peaks. The number in parentheses denotes the count of genes with TSS located within 10 kb of the significantly differential peaks in each module. NES, normalized enrichment score.

**Figure supplement 6.**
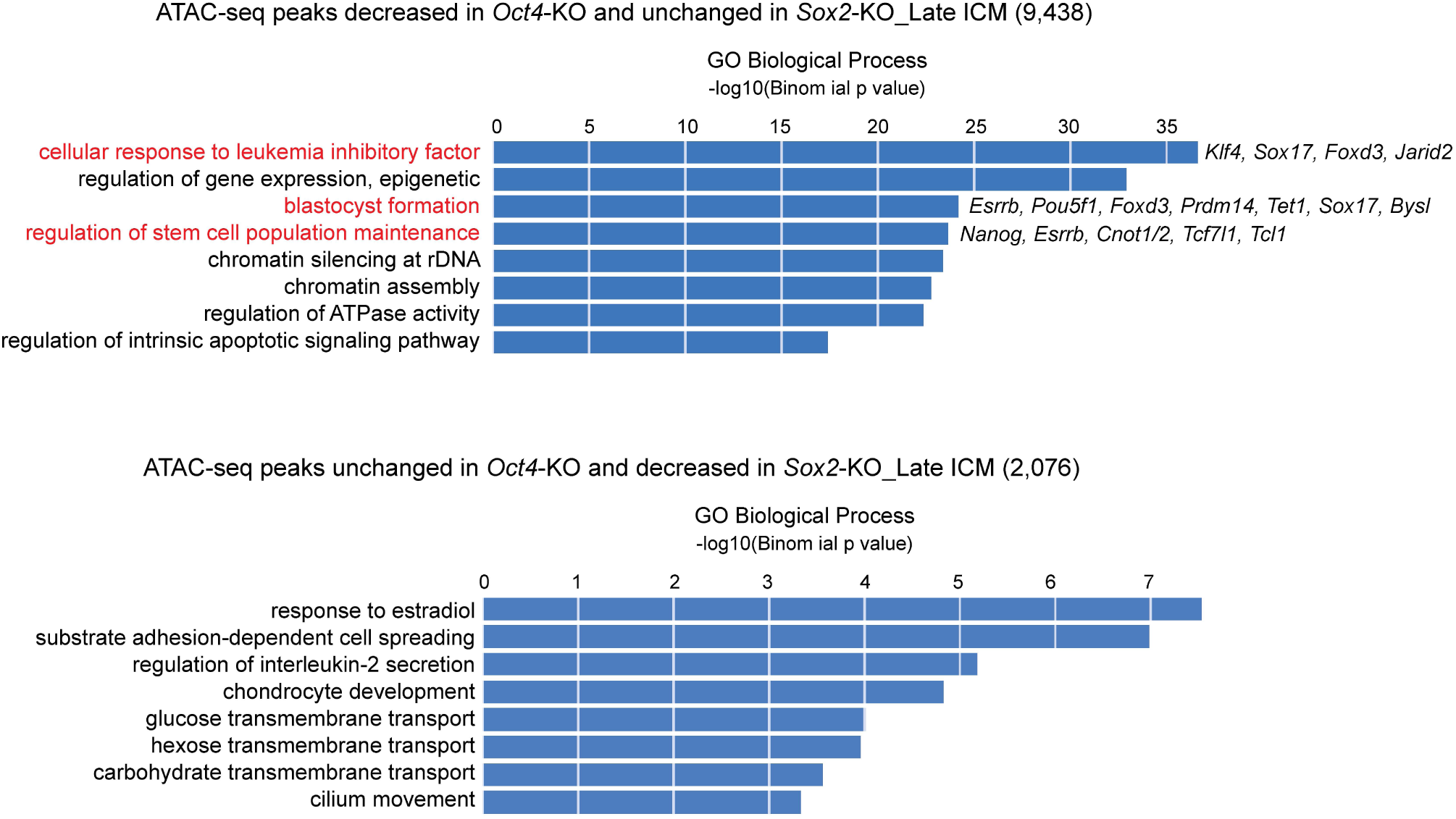
GREAT Biological Process enrichment analysis of ATAC-seq peaks specifically decreased in the *Oct4-*KO or *Sox2-*KO late ICMs (related to Figure 3) Terms highlighted in red are related to the pluripotency and preimplantation embryonic development. Example genes of associated peaks are listed.

**Figure supplement 7.**
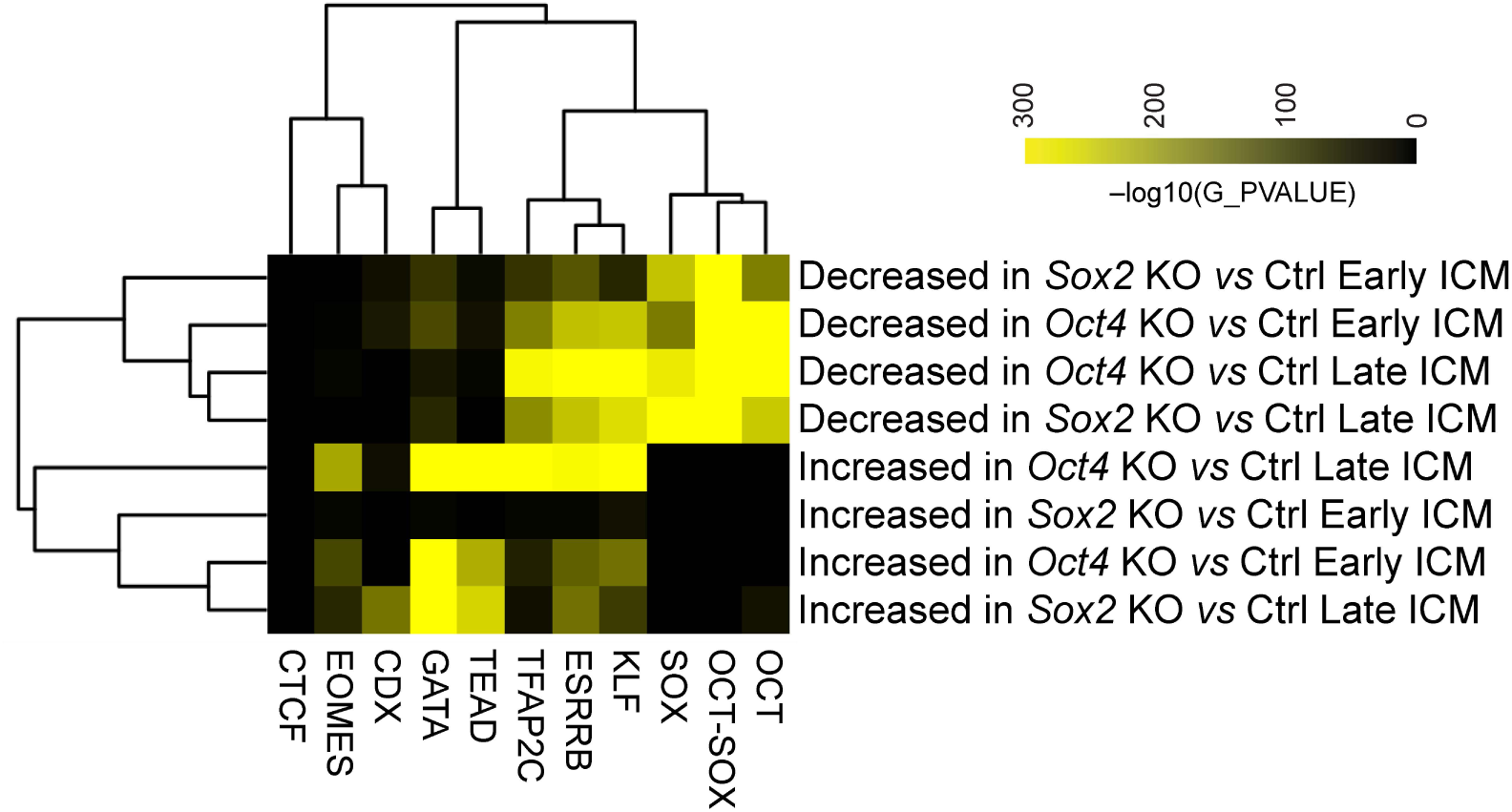
Motif enrichment analysis of significantly changed ATAC-seq peaks that are distal to TSSs (related to Figure 4). The values are –log_10_(p-value)

**Figure supplement 8.**
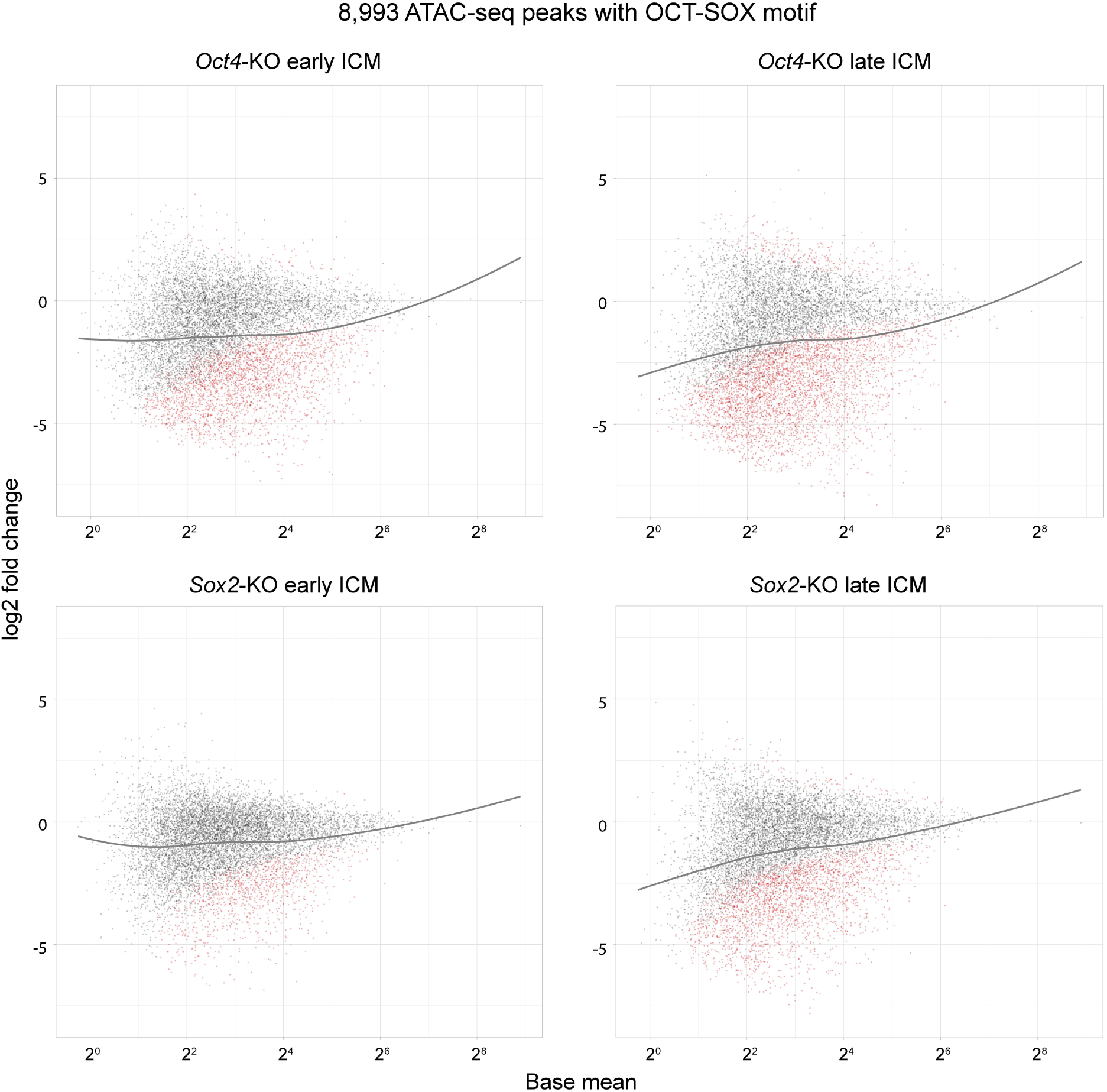
MA plots show the log_2_ fold change of the 8,993 OCT-SOX peaks in *Oct4*- and *Sox2*-KO ICMs (related to Figure 4) Black and red dots represent the unchanged and significantly changed peaks, respectively. Grey lines are the regression curves of the 8,993 OCT-SOX peaks.

**Figure supplement 9.**
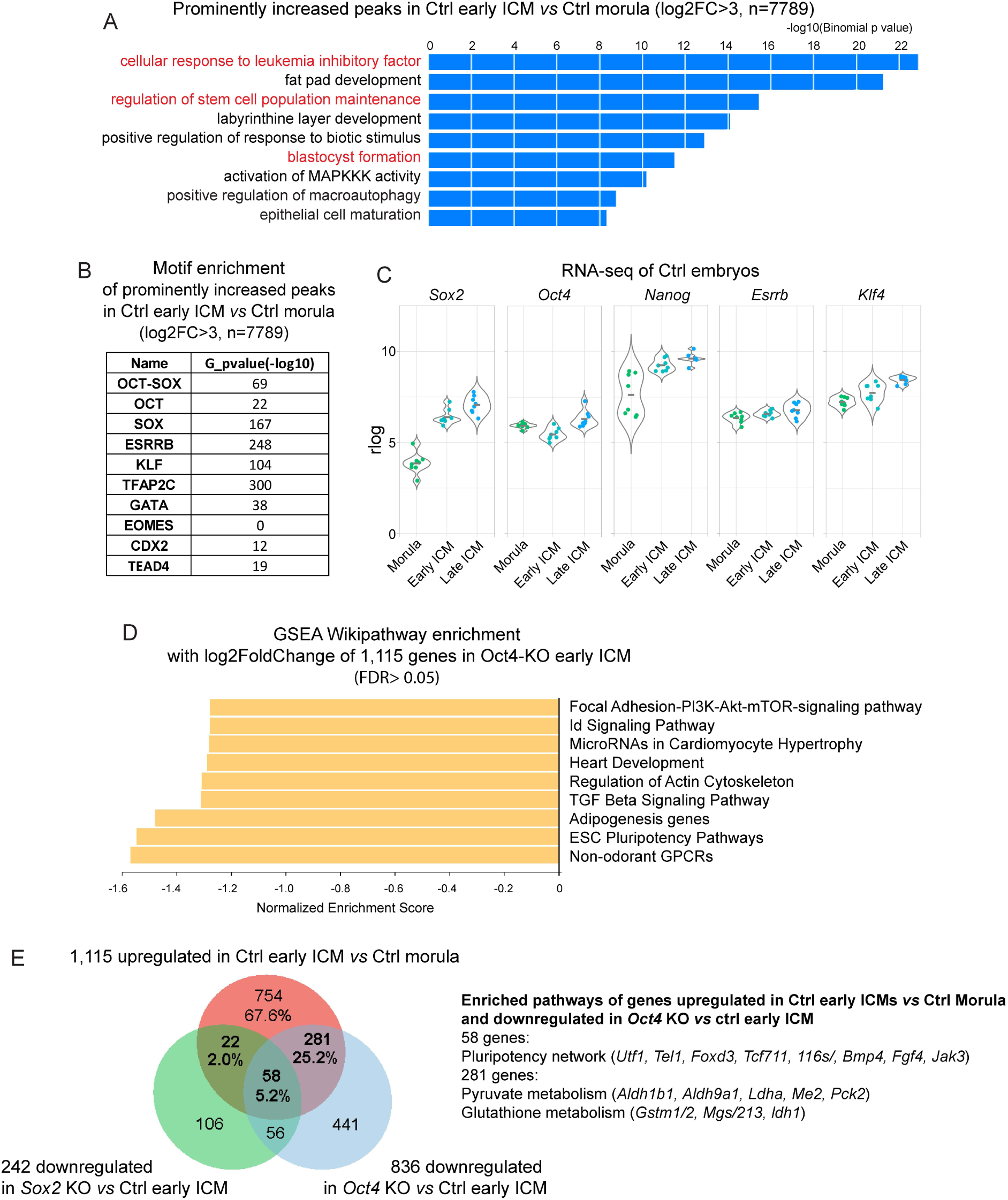
The dynamics of chromatin accessibility and transcriptome from morula to ICM (related to Figure 5) **A** GREAT analysis was conducted on the most prominently elevated ATAC-seq peaks in *Ctrl* early ICM *vs Ctrl* morula. A log2 fold change cutoff of >3 was applied due to the limits of GREAT analysis with fewer than 10,000 peaks. **B**. Motif enrichment analysis with PScan of the increased ATAC-seq peaks in *Ctrl* early ICM *vs Ctrl* morula. Cutoff, log2 fold change >3. C. Expression of *Sox2*, *Oct4*, *Nanog*, *Esrrb* and *Klf4* in morulae and ICMs (related to Figure 5). **D.** Bar chart illustrating the GSEA Wikipathway enrichment in WebGestalt. The log2 foldchange values of the 1,115 upregulated genes (Figure 5D) in *Oct4* KO *vs* Ctrl early ICM were used in this analysis. FDR >0.05. **E.** Venn diagram showing the overlap between genes upregulated in early ICMs and those downregulated in *Oct4-*KO or *Sox2-*KO early ICMs. Pathway over-representation analysis was conducted on the genes upregulated in Ctrl early ICMs *vs* Ctrl Morula and downregulated in *Oct4* KO *vs* Ctrl early ICM.

