## Supplementary material for "Emerging cooperativity between Oct4 and Sox2 governs the pluripotency network in early mouse embryos": Table supplement 1. Coordinates for the marked well-known enhancers in Figure 3A

**Table supplement 1 Coordinates for marked enhancers in Fig. 3A**

| **Gene of the enhancer** | **Chr** | **start** | **end** | **coordinate** |
| --- | --- | --- | --- | --- |
| Prdm14 | chr1 | 13127092 | 13127292 | chr1:13127154-13127231 |
| Pecam1 | chr11 | 106724458 | 106724658 | chr11:106724438-106724678 |
| Pou5f1_CR4 | chr17 | 35503917 | 35504117 | chr17:35503933-35504102 |
| Sox2_SRR2 | chr3 | 34653964 | 34654164 | chr3:34654015-34654114 |
| Klf4-B | chr4 | 55477518 | 55477718 | chr4:55477392-55477844 |
| Nanog_DE | chr6 | 122702564 | 122702764 | chr6:122702605-122702724 |
| Nanog_PE | chr6 | 122707419 | 122707619 | chr6:122707461-122707577 |
| Utf1 | chr7 | 139945619 | 139945819 | chr7:139945645-139945793 |
| Klf2 | chr8 | 72334360 | 72334560 | chr8:72334319-72334601 |
