## Supplementary material for "Emerging cooperativity between Oct4 and Sox2 governs the pluripotency network in early mouse embryos": Table supplement 2. oligos for ATAC-seq library preparation

**Table supplement 2. The sequences of the indexes used in ATAC-seq library preparation.**

| **Group** | **Primer name** | **Sequence** | **Index** |
| --- | --- | --- | --- |
| Index_1 | Index_1.1 (N701) | CAAGCAGAAGACGGCATACGAGATTCGCCTTAGTCTCGTGGGCTCGGAGATGT | TCGCCTTA |
| Index_1 | Index_1.2 (N702) | CAAGCAGAAGACGGCATACGAGATCTAGTACGGTCTCGTGGGCTCGGAGATGT | CTAGTACG |
| Index_1 | Index_1.3 (N703) | CAAGCAGAAGACGGCATACGAGATTTCTGCCTGTCTCGTGGGCTCGGAGATGT | TTCTGCCT |
| Index_1 | Index_1.4 (N704) | CAAGCAGAAGACGGCATACGAGATGCTCAGGAGTCTCGTGGGCTCGGAGATGT | GCTCAGGA |
| Index_1 | Index_1.5 (N705) | CAAGCAGAAGACGGCATACGAGATAGGAGTCCGTCTCGTGGGCTCGGAGATGT | AGGAGTCC |
| Index_1 | Index_1.6 (N706) | CAAGCAGAAGACGGCATACGAGATCATGCCTAGTCTCGTGGGCTCGGAGATGT | CATGCCTA |
| Index_2 | Index_2.1 (501) | AATGATACGGCGACCACCGAGATCTACACTAGATCGCTCGTCGGCAGCGTCAGATGTG | TAGATCGC |
| Index_2 | Index_2.2 (502) | AATGATACGGCGACCACCGAGATCTACACCTCTCTATTCGTCGGCAGCGTCAGATGTG | CTCTCTAT |
| Index_2 | Index_2.3 (503) | AATGATACGGCGACCACCGAGATCTACACTATCCTCTTCGTCGGCAGCGTCAGATGTG | TATCCTCT |
| Index_2 | Index_2.4 (504) | AATGATACGGCGACCACCGAGATCTACACAGAGTAGATCGTCGGCAGCGTCAGATGTG | AGAGTAGA |
| Index_2 | Index_2.5 (505) | AATGATACGGCGACCACCGAGATCTACACGTAAGGAGTCGTCGGCAGCGTCAGATGTG | GTAAGGAG |
| Index_2 | Index_2.6 (506) | AATGATACGGCGACCACCGAGATCTACACACTGCATATCGTCGGCAGCGTCAGATGTG | ACTGCATA |
| Index_2 | Index_2.7 (507) | AATGATACGGCGACCACCGAGATCTACACAAGGAGTATCGTCGGCAGCGTCAGATGTG | AAGGAGTA |
| Index_2 | Index_2.8 (508) | AATGATACGGCGACCACCGAGATCTACACCTAAGCCTTCGTCGGCAGCGTCAGATGTG | CTAAGCCT |
