## Supplementary material for "Emerging cooperativity between Oct4 and Sox2 governs the pluripotency network in early mouse embryos": Table supplement 3. oligos for RNA-seq library preparation

**Table supplement 3. The sequences of the oligos and indexes used in RNA-seq library preparation.**

| **Name** | **Usage** | **Sequence** |
| --- | --- | --- |
| R2SP-UMI-(dT)24 | Reverse transcription | GTGACTGGAGTTCAGACGTGTGCTCTTCCGATCTNNNNNNNNNNTTTTTTTTTTTTTTTTTTTTTTTT |
| R2SP | 1st amplification | GTGACTGGAGTTCAGACGTGTGCTCTTCCGATCT |
| V3(dT)24 | 1st amplification | ATATCTCGAGGGCGCGCCGGATCCTTTTTTTTTTTTTTTTTTTTTTTT |
| NH2-R2SP | 2nd amplification | NH2-GTGACTGGAGTTCAGACGTGTGCTCTTCCGATCT |
| NH2-V3(dT)24 | 2nd amplification | NH2-ATATCTCGAGGGCGCGCCGGATCCTTTTTTTTTTTTTTTTTTTTTTTT |
| R1SP | Adaptor | ACACTCTTTCCCTACACGACGCTCTTCCGATC*T |
| R1SP_Complement | Adaptor | GATCGGAAGAGC |
| i701-R2SP | Index | CAAGCAGAAGACGGCATACGAGATCGAGTAATGTGACTGGAGTTCAGACGTGTGCTCTTCCGATC*T |
| i702-R2SP | Index | CAAGCAGAAGACGGCATACGAGATTCTCCGGAGTGACTGGAGTTCAGACGTGTGCTCTTCCGATC*T |
| i703-R2SP | Index | CAAGCAGAAGACGGCATACGAGATAATGAGCGGTGACTGGAGTTCAGACGTGTGCTCTTCCGATC*T |
| i704-R2SP | Index | CAAGCAGAAGACGGCATACGAGATGGAATCTCGTGACTGGAGTTCAGACGTGTGCTCTTCCGATC*T |
| i705-R2SP | Index | CAAGCAGAAGACGGCATACGAGATTTCTGAATGTGACTGGAGTTCAGACGTGTGCTCTTCCGATC*T |
| i706-R2SP | Index | CAAGCAGAAGACGGCATACGAGATACGAATTCGTGACTGGAGTTCAGACGTGTGCTCTTCCGATC*T |
| i707-R2SP | Index | CAAGCAGAAGACGGCATACGAGATAGCTTCAGGTGACTGGAGTTCAGACGTGTGCTCTTCCGATC*T |
| i708-R2SP | Index | CAAGCAGAAGACGGCATACGAGATGCGCATTAGTGACTGGAGTTCAGACGTGTGCTCTTCCGATC*T |
| i709-R2SP | Index | CAAGCAGAAGACGGCATACGAGATCATAGCCGGTGACTGGAGTTCAGACGTGTGCTCTTCCGATC*T |
| i710-R2SP | Index | CAAGCAGAAGACGGCATACGAGATTTCGCGGAGTGACTGGAGTTCAGACGTGTGCTCTTCCGATC*T |
| i711-R2SP | Index | CAAGCAGAAGACGGCATACGAGATGCGCGAGAGTGACTGGAGTTCAGACGTGTGCTCTTCCGATC*T |
| i712-R2SP | Index | CAAGCAGAAGACGGCATACGAGATCTATCGCTGTGACTGGAGTTCAGACGTGTGCTCTTCCGATC*T |
| i713-R2SP | Index | CAAGCAGAAGACGGCATACGAGATAGGAGGAAGTGACTGGAGTTCAGACGTGTGCTCTTCCGATC*T |
| i714-R2SP | Index | CAAGCAGAAGACGGCATACGAGATAGCAAGCAGTGACTGGAGTTCAGACGTGTGCTCTTCCGATC*T |
| i715-R2SP | Index | CAAGCAGAAGACGGCATACGAGATTCATCACCGTGACTGGAGTTCAGACGTGTGCTCTTCCGATC*T |
| i716-R2SP | Index | CAAGCAGAAGACGGCATACGAGATCGTAGGTTGTGACTGGAGTTCAGACGTGTGCTCTTCCGATC*T |
| i717-R2SP | Index | CAAGCAGAAGACGGCATACGAGATTCAGATCCGTGACTGGAGTTCAGACGTGTGCTCTTCCGATC*T |
| i718-R2SP | Index | CAAGCAGAAGACGGCATACGAGATCGTGATCAGTGACTGGAGTTCAGACGTGTGCTCTTCCGATC*T |
| i719-R2SP | Index | CAAGCAGAAGACGGCATACGAGATAGTCGCTTGTGACTGGAGTTCAGACGTGTGCTCTTCCGATC*T |
| i720-R2SP | Index | CAAGCAGAAGACGGCATACGAGATGAACGCTTGTGACTGGAGTTCAGACGTGTGCTCTTCCGATC*T |
| i721-R2SP | Index | CAAGCAGAAGACGGCATACGAGATTACGCCTTGTGACTGGAGTTCAGACGTGTGCTCTTCCGATC*T |
| i722-R2SP | Index | CAAGCAGAAGACGGCATACGAGATCTCATCAGGTGACTGGAGTTCAGACGTGTGCTCTTCCGATC*T |
| i723-R2SP | Index | CAAGCAGAAGACGGCATACGAGATTCTTCTGCGTGACTGGAGTTCAGACGTGTGCTCTTCCGATC*T |
| i724-R2SP | Index | CAAGCAGAAGACGGCATACGAGATGCTGGATTGTGACTGGAGTTCAGACGTGTGCTCTTCCGATC*T |
| i501 | Index | AATGATACGGCGACCACCGAGATCTACACTATAGCCTACACTCTTTCCCTACACGACGCTCTTCCGATC*T |
| i502 | Index | AATGATACGGCGACCACCGAGATCTACACATAGAGGCACACTCTTTCCCTACACGACGCTCTTCCGATC*T |
| i503 | Index | AATGATACGGCGACCACCGAGATCTACACCCTATCCTACACTCTTTCCCTACACGACGCTCTTCCGATC*T |
| i504 | Index | AATGATACGGCGACCACCGAGATCTACACGGCTCTGAACACTCTTTCCCTACACGACGCTCTTCCGATC*T |
| i505 | Index | AATGATACGGCGACCACCGAGATCTACACAGGCGAAGACACTCTTTCCCTACACGACGCTCTTCCGATC*T |
| i506 | Index | AATGATACGGCGACCACCGAGATCTACACTAATCTTAACACTCTTTCCCTACACGACGCTCTTCCGATC*T |
| i507 | Index | AATGATACGGCGACCACCGAGATCTACACCAGGACGTACACTCTTTCCCTACACGACGCTCTTCCGATC*T |
| i508 | Index | AATGATACGGCGACCACCGAGATCTACACGTACTGACACACTCTTTCCCTACACGACGCTCTTCCGATC*T |
| i509 | Index | AATGATACGGCGACCACCGAGATCTACACTTGCTTGCACACTCTTTCCCTACACGACGCTCTTCCGATC*T |
| i510 | Index | AATGATACGGCGACCACCGAGATCTACACGAGAGGTTACACTCTTTCCCTACACGACGCTCTTCCGATC*T |
| i511 | Index | AATGATACGGCGACCACCGAGATCTACACACCTGGTTACACTCTTTCCCTACACGACGCTCTTCCGATC*T |
| i512 | Index | AATGATACGGCGACCACCGAGATCTACACAAGCGGAAACACTCTTTCCCTACACGACGCTCTTCCGATC*T |
| i513 | Index | AATGATACGGCGACCACCGAGATCTACACCGGAACAAACACTCTTTCCCTACACGACGCTCTTCCGATC*T |
| i514 | Index | AATGATACGGCGACCACCGAGATCTACACGGTAAGCTACACTCTTTCCCTACACGACGCTCTTCCGATC*T |
| i515 | Index | AATGATACGGCGACCACCGAGATCTACACTGTGGCATACACTCTTTCCCTACACGACGCTCTTCCGATC*T |
| i516 | Index | AATGATACGGCGACCACCGAGATCTACACACTACGGAACACTCTTTCCCTACACGACGCTCTTCCGATC*T |
